# Platelet-derived LPA16:0 inhibits adult neurogenesis and stress resilience in anxiety disorder

**DOI:** 10.1101/2024.08.18.608479

**Authors:** Thomas Larrieu, Charline Carron, Fabio Grieco, Crystal Weber, Kyllian Ginggen, Aurélie Delacrétaz, Hector Gallart-Ayala, Mumeko Tsuda, Heather A. Cameron, Chin B. Eap, Julijana Ivanisevic, Pierre Magistretti, Ludovic Telley, Alexandre Dayer, Camille Piguet, Nicolas Toni

## Abstract

Anxiety disorders are accompanied by changes in brain plasticity, stress vulnerability and heightened risk of depression. Here, we found that serum LPA16:0 abundance increased with trait anxiety in both human and mice and was sufficient to reduce the proliferation of adult hippocampal neural stem/progenitor cells. In humans, the main LPA receptor, LPA_1_, bears single nucleotide polymorphism variants associated with anxiety. In mice, LPA16:0 decreased hippocampal neurogenesis and stress resilience, whereas LPA_1_ antagonism or the reduction of platelets, the main source of circulating LPA16:0, increased adult neurogenesis and resilience to acute stress. Finally, the inhibition of adult neurogenesis abolished the beneficial effect of LPA_1_ antagonism on resilience against both acute and chronic stress.

Together, these findings identify LPA16:0-LPA_1_ signaling as a regulation mechanism of adult neurogenesis and a potential therapeutic target for mood disorders.

## Introduction

Anxiety disorders represent the most prevalent forms of mental illnesses, affecting 20-30% of the population aged 15-25 years^1^. Their high chronicity and prevalence contribute to their substantial impact on global health, accounting for approximately 3% of the overall burden of disease^2^. Moreover, anxiety frequently co-occurs with various psychiatric conditions, exacerbating the severity of these disorders and increasing the risk of recurrence or relapse^3–6^. Unfortunately, anxiety often goes untreated, leading to prolonged suffering for individuals and a persistent burden on healthcare systems^7–9^. Thus, to gain a deeper understanding of the underlying mechanisms driving anxiety and its interplay with other psychiatric disorders, adopting a trans-diagnostic framework that incorporates anxiety as a fundamental dimension is crucial. Implementing a trans-diagnostic perspective can also pave the way for personalized treatment approaches.

Individuals diagnosed with anxiety disorders often exhibit heightened reactivity to stress^10^. Pathological stress response involves alterations in brain plasticity^11,12^, in particular impacting adult hippocampal neurogenesis^13^. Adult neurogenesis results in the continuous generation of new neurons in the adult brain, a process observed in the dentate gyrus (DG) of the hippocampus^14^. Interestingly, adult neurogenesis is highly sensitive to changes in lifestyle conditions such as social isolation^15^, inflammation^16,17^ or chronic stress^17–20^, all of which are associated with decreased neurogenesis and emotional deficits^11,16,21^. Conversely, the production of new granule neurons is crucial for regulating mood^20,22,23^, promoting stress resilience^24–26^, and mediating the effects of antidepressant treatments^20,27^ and increasing adult neurogenesis itself can produce antidepressant effects^28^. Thus, adult neurogenesis is both a target of anxiety and stress-related disorders as well as a mediator of stress resilience. However, the mechanisms underlying the effect of anxiety on stress vulnerability and adult neurogenesis are not fully understood. Mechanistically, most of the effects of external factors on adult neurogenesis are mediated by the close cellular environment of aNPC, named the neurogenic niche, which includes neurons, astrocytes, oligodendrocytes, microglia, as well as blood vessels. The concept of the vascular neurogenic niche was initially based on observations of the proximity between hippocampal stem/progenitor cells and blood vasculature^29^ where capillaries serve as a scaffold for progenitor cells’ migration^30^. Ultrastructural observations further showed that hippocampal radial glia-like stem cells form perivascular endfeet at sites of increased capillaries permeability^31,32^, suggesting that blood-circulating molecules may directly interact with stem cells and regulate their proliferation and fate despite poor blood-brain barrier permeability. In support of this possibility, several studies have uncovered several blood-borne molecules that regulate adult neurogenesis in age-related cognitive impairment or voluntary exercise^33–38^. However, the role of blood-circulating molecules in the regulation of adult neurogenesis in the context of stress and anxiety is poorly known.

## Results

### Serum from anxious mice reduces aNPC proliferation

To investigate blood-circulating molecules that regulate adult hippocampal neurogenesis in the context of anxiety, we developed an *in vitro* model in which adult hippocampal neural stem/progenitor cells (aNPC) were exposed to serum from naïve adult C57BL6/J male mice. aNPC were cultured in presence of serum for either 24 hours under adherent conditions or 72 hours in floating conditions and their proliferation was assessed using the thymidine analogue bromodeoxyuridine (BrdU) and a neurosphere assay (Fig. 1A). In both conditions, increasing serum concentrations decreased aNPC proliferation (Fig. 1B-E). However, a concentration of 0.2% serum did not affect proliferation as compared to serum-free culture medium. We therefore kept this concentration and the adherent cell culture condition for all further experiments and named this assay the blood-brain axis (BBA) assay. We next used the BBA assay to examine the effect of serum sampled from inbred adult C57BL/6J male mice under physiological (i.e., stress-free and undisturbed housing) conditions. Under these conditions, rodents exhibit natural variations in anxiety-related behavior^39,40^, which are associated with opposite variations of adult neurogenesis^41^. Mice were first monitored for baseline inter-individual differences in anxiety (i.e., trait-anxiety) using both an elevated plus maze (EPM) and a light/dark test (LDT) (Fig. 1F, G), and results from these tests were then integrated into a trait-anxiety score that results from the average of the normalized score on the two tests (see Materials and Methods). Mice displayed a great variability in their anxiety score (Fig. 1H) and the group was subsequently divided along the median into low anxious (LA) and high anxious mice (HA) (Fig. 1I). Eight days after behavioral testing serum from each mouse was collected and tested in the BBA assay. aNPC proliferation was reduced upon exposure to the serum obtained from HA mice, as compared to the serum from LA mice (Fig. 1J). Furthermore, aNPC proliferation in the BBA assay negatively correlated with individual trait-anxiety scores (Fig. 1K), indicating that aNPC proliferation was inhibited by blood-circulating molecules that vary according to the basal emotional status of each individual. To bolster the validity of these findings, the experiment was repeated using a separate group of naïve mice. In this independent cohort, we observed a 15% decrease in the proliferation of adult neural progenitor cells (aNPCs) when they were exposed to serum obtained from high-anxiety (HA) mice compared to low-anxiety (LA) mice (LA: 100 ± 4.57%, n=11; HA: 84.87 ± 4.67%, n=9; p=0.034). Since glucocorticoids exert a bimodal, dose-dependent effect on aNPC proliferation *in vitro*^42^, we measured the free corticosterone (CORT) levels in the serum of low- and high-anxious mice. Free-CORT levels were similar between HA and LA mice (Fig. 1L) and did not correlate with aNPC proliferation in the BBA assay (Fig. 1M). Similarly, to test the possibility that serum toxicity may also account for a decrease in cell proliferation, we examined nuclear condensation, a morphological cardinal feature of apoptosis^43^, as well as caspase 3 activation. We did not find any difference between groups for nuclear condensation (Supp. Fig. 1A, B) or caspase 3 activation (Supp. Fig. 1C, D), indicating that serum from HA mice did not induce toxicity *in vitro*. We next examined the relationship between anxiety and adult neurogenesis *in vivo*. To this aim, the mice were injected with BrdU one day prior to blood sampling and perfusion and hippocampal sections were immunostained for BrdU (Fig. 1F). The number of BrdU-immunolabeled cells was reduced in the dentate gyrus of HA compared to LA mice (Supp. Fig. 2A). Furthermore, the number of BrdU-immunolabeled cells negatively correlated with the trait-anxiety score (Supp. Fig. 2B). We also examined the relative abundance of the immature neuron marker doublecortin (DCX) and we found a nonsignificant decrease in doublecortin (DCX) staining in the DG of HA compared to LA mice (Supp. Fig. 2C) and no correlation with trait-anxiety (Supp. Fig. 2D). Together, these results indicate that the BBA assay enables the examination of blood circulating molecules that inhibit adult hippocampal neurogenesis in the context of basal (trait) anxiety.

**Fig. 1:**
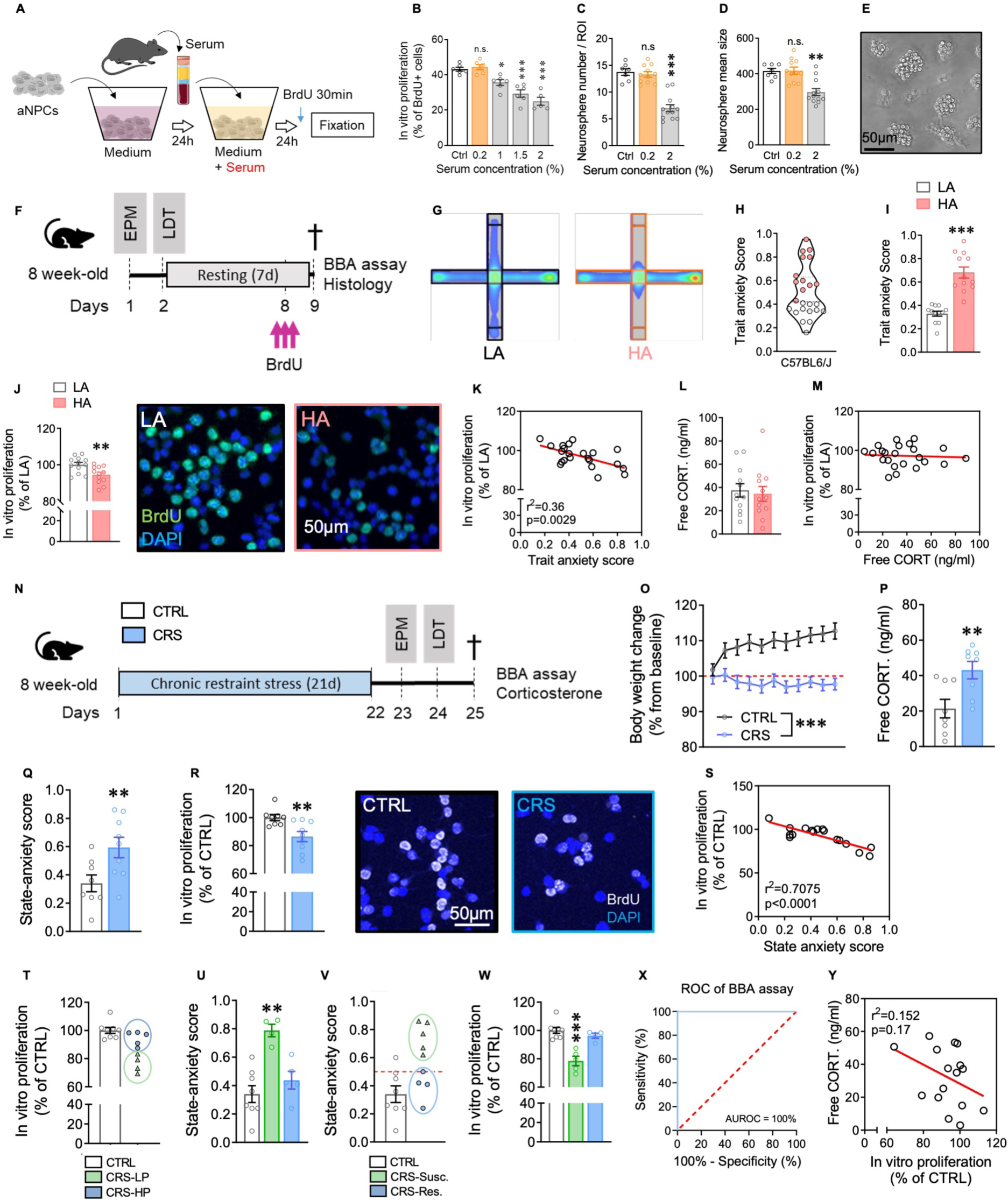
Serum from anxious mice reduces aNPC proliferation *in vitro*. A. Schematic illustration of the BBA assay. **B**. Proportion of aNPC that incorporated BrdU after exposure to serum from 4 adult C57BL/6J male mice at different concentrations for 24 hours under coating conditions. F4,24 = 23.32, p < 0.0001, one-way ANOVA; Tukey’s multiple comparisons with the mean of CTRL group, 2 wells per condition, 3 independent replicates. **C, D.** Number (**C**) and mean size (**D**) of neurospheres after exposure to serum from 4 adult C57BL/6J mice at different concentrations for 72 hours. Neurospheres number: F2,27 = 40.12, p < 0.0001, one-way ANOVA; Tukey’s multiple comparisons with the mean of CTRL group; Neurospheres size: F2,27 = 13.55, p < 0.0001, one-way ANOVA; Tukey’s multiple comparisons with the mean of CTRL group. The number and mean size of neurospheres were counted in 4 selected fields per well, 3 wells per condition. **E.** Representative bright field image of neurospheres. **F.** Experimental design for the anxiety assessment. **G.** Grouped heatmap of the time spent in the open (vertical) and closed (horizontal) arms of an EPM for low (LA) and high (HA) anxious mice. **H-I.** Violin plot (**H**) and histogram (**I**) of the anxiety score of LA (white) and HA (red) mice. **I.** t22 = 6.79, p < 0.0001, unpaired t-test, two-tailed, n = 12 per group. **J (left)**. Histogram of aNPC proliferation in the BBA assay after exposure to 0.2 % of serum from high or low anxious mice (t22 = 2.55, p = 0.018, unpaired t-test, two-tailed, n = 12 per group). **J (right):** Representative confocal micrographs of aNPC immunostained for BrdU and DAPI after exposure to serum from LA and HA mice. **K**. Correlation plot between *in vitro* proliferation and trait anxiety score. **L**. Histogram of the free serum corticosterone in LA and HA mice (t22 = 0,3519, p > 0.05 unpaired t-test, two-tailed, n = 12 per group). **M**. Correlation plot between *in vitro* cell proliferation and free corticosterone. **N**. Experimental design for the chronic restraint stress (CRS) experiment. n=8 CRS mice and n=9 non-stressed control (CTRL) mice. **O**. Body weight change over time (Stress effect: F(1, 15) = 16,61; p = 0,001, Two-way ANOVA with repeated measure). **P**. Free serum corticosterone (t14 = 3,021, p = 0.009, unpaired t-test, two-tailed, n = 8 per group). **Q**. State anxiety score (t15 = 3.08, p = 0.0082, unpaired t-test, two-tailed, n = 8 and 9 per group). **R (left)**. Cell proliferation in the BBA assay after treatment with serum from CRS or CTRL mice (t15 = 3.11, p = 0.0067, unpaired t-test, two-tailed, n = 8 and 9 per group). **R (right)**. Representative confocal micrographs of aNPC treated with serum from CTRL or CRS mice and immunostained for BrdU (white) and DAPI (blue). **S**. Correlation plot between cell proliferation in the BBA assay and the state anxiety score. **T**. Same data as in panel **R**, where CRS-treated mice were subdivided in high proliferative (HP, blue) and low proliferative (LP, green). **U**. State anxiety score of CTRL, CRS-HP and CRS-LP mice. (p = 0.0030, Kruskal-Wallis test, post-hoc analysis for multiple comparisons: CTRL vs. CRS-LP, p = 0.0047). **V**. Same data as in panel **Q**, where CRS-treated mice were subdivided in resilient (CRS-res, blue) and susceptible (CRS-susc, green) mice. **W**. Cell proliferation in the BBA assay of CTRL, CRS-resilient and CRS-susceptible mice (p = 0.006, Kruskal-Wallis test, post-hoc analysis for multiple comparisons: CTRL vs. CRS-Susc., p = 0.0034). **X**. Receiver Operating Characteristic (ROC) curve for the discrimination of CRS-induced susceptibility by the BBA assay. **Y**. Correlation between proliferation in the BBA assay and free serum corticosterone. For correlation plots, linear regression, r^2^ and p values are indicated in the graphs when significant correlations were found. Histograms show average ± SEM. ns: not significant; * p<0.05; ** p<0.01; *** p<0.001.

### The BBA assay identifies stress-resilient individuals

We next assessed the BBA assay in the context of state anxiety induced by chronic stress. To this aim, we subjected mice to a 21-day-chronic restraint stress (CRS) regimen, which induces anxiety and depression-like symptoms^11,44,45^. We then assessed their anxiety scores and, 24 hours later, we sampled their sera for assessment in the BBA assay (Fig. 1N). As expected, CRS-treated mice displayed no weight gain (Fig. 1O), increased free CORT serum concentration (Fig. 1P) and increased state-anxiety scores (Fig. 1Q) as compared to non-stressed mice, thus confirming the effectiveness of the CRS protocol. We then assessed the sera in the BBA assay and found that serum from CRS-treated mice reduced aNPC proliferation as compared to serum from non-stressed mice (Fig. 1R). We replicated this experiment in an independent cohort of mice, in which we found a 14% reduction of aNPC proliferation in CRS-treated mice compared with control mice (data not shown, CTRL, 100 ± 4.36%, n=11; CRS, 86.02 ± 4.53%, n=11; p=0.037). Furthermore, aNPC proliferation inversely correlated with the state-anxiety score following CRS (Fig. 1S), indicating that the BBA assay was sensitive to blood circulating molecules related to stress-induced anxiety. Interestingly, when CRS-treated mice were divided along the median into low-proliferative and high-proliferative mice (Fig. 1T), only mice from which the serum inhibited cell proliferation in the BBA assay displayed a high anxiety score (Fig. 1U), suggesting that the BBA assay detected a stress-sensitive phenotype. Inversely, when CRS-treated mice were divided along the median of the anxiety score (Fig. 1V), only mice in the high-anxiety group displayed reduced cell proliferation in the BBA assay (Fig. 1W), suggesting that the BBA assay discriminated susceptible from resilient animals. Subsequently, we plotted the true positive values (Specificity) against the false positive values (Sensitivity) using the Receiver Operating Characteristic (ROC) curve. The BBA assay showed a 100% accuracy at discriminating between resilient and susceptible animals after CRS (Fig. 1X). Although we observed an increase in free-CORT levels in CRS-treated mice, we found no significant correlation between free-CORT levels and *in vitro* proliferation in the BBA assay (Fig. 1Y), indicating that corticosterone did not mediate the effect of serum on aNPC proliferation in the BBA assay under these conditions. We also excluded any cytotoxicity effect of serum from CRS-treated mice on aNPC proliferation in the BBA assay using the quantification of nuclear condensation and immunocytochemistry for Caspase 3 activation (Supp. Fig. 1E-H). Together, these results indicate that blood-circulating molecules regulate aNPC proliferation in the context of state-anxiety induced by experimental stress.

### LPA16:0-LPA_1_ signaling is involved in anxiety in patients

We next aimed to test human serum on the BBA assay. First, we tested the compatibility of the BBA assay with human serum by assessing the serum of 5 healthy control individuals. The concentration of 0.2% serum from human controls did not affect aNPC proliferation as compared to serum from control mice, (data not shown. Mouse serum, 35.16 ± 1.49%, n=11; Human serum, 38.30 ± 0.88%, n=5; p=0.20), indicating that the BBA assay is applicable to human serum. We then examined the effect of serum from individuals from the BipOff cohort of the Geneva University Hospital, which consists in young adults with at least one parent with a history of neuropsychiatric disorder related to emotional dysregulation (Attention-Deficit Hyperactivity Disorder, Borderline Personality Disorder, or Bipolar Disorder). These young individuals are known to have increased risk of developing a concordant or a non-concordant psychiatric disorder^46–52^. Although the study was cross-sectional, at the time of blood sampling, some individuals had developed a neuropsychiatric disorder, and some had remained healthy. They were therefore categorized as high-risk (HR)-susceptible or HR-resilient respectively (see Materials and Methods and Fig. 2A, B). HR-susceptible individuals displayed higher scores of trait- and state-anxiety, as measured by self-reported questionnaires, as compared to both healthy and HR-resilient subjects (Fig. 2C, D). In addition, the serum from HR-susceptible individuals significantly decreased aNPC proliferation in the BBA assay, as compared to serum from healthy controls or HR-resilient individuals which displayed comparable proliferation (Fig. 2E, F). The area under the ROC curve indicated a 70% accuracy at discriminating HR-susceptible individuals from healthy controls (Fig. 2G). Further analysis of high-risk individuals revealed an inverse correlation between aNPCs proliferation in the BBA assay and anxiety (trait and state) and individuals with the greatest reduction of cell proliferation in the BBA assay displayed the highest clinical scores for both trait- and state-anxiety (Fig. 2H, I). Importantly, we did not observe a correlation between cortisol levels and cell proliferation in the BBA assay (data not shown, r^2^=0.038, p=0.28), suggesting that other circulating molecules related to anxiety regulate aNPC proliferation in the BBA assay. Moreover, serum from HR individuals did not influence cell viability compared to healthy serum (Supp. Fig. 3A, B). Overall, we found remarkable similarities between mouse and human serum, indicating that blood-circulating molecules associated with anxiety mediated the reduction of aNPC proliferation in the BBA assay.

**Fig. 2.**
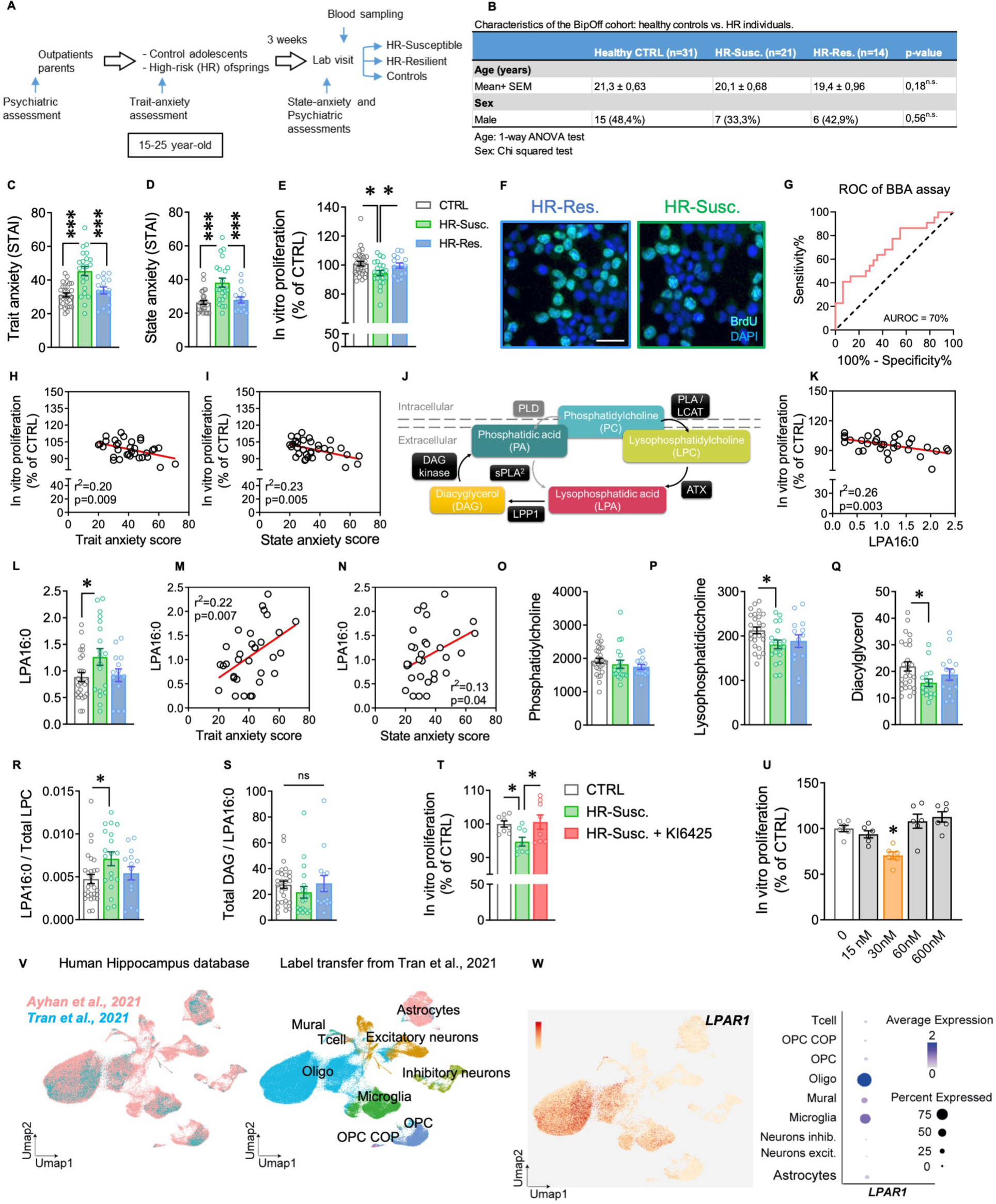
LPA16:0 increases with anxiety in patients and induces a LP1-dependent reduction in aNSC proliferation. A. Schematic illustration of patients’ assessment. **B**. Characteristics of the BipOff cohort. **C, D.** Trait (**C**) and state (**D**) anxiety levels in CTRL, HR-S and HR-Res individuals (**C,** F2,65 = 15.16, p < 0.0001, one-way ANOVA, **D**, F2,65 = 11.83, p < 0.0001, one-way ANOVA). **E**. Cell proliferation in the BBA assay after treatment with serum from CTRL, HR-S and HR-Res individuals (F2,65 = 4.06, p = 0.022, one-way ANOVA). **F**. Representative confocal micrographs of aNPC treated with serum from HR-S and HR-Res individuals and immunostained against BrdU (green) and DAPI (blue). **G**. Receiver Operating Characteristic (ROC) curve fort the discrimination of HR-S individuals by the BBA assay. **H, I**. Correlations between cell proliferation in the BBA assay and trait (**H**) or state (**I**) anxiety. **J**. Schematic illustration of the biosynthesis and degradation of LPA (Phospholipase D, PLD; Phospholipase A, PLA; Diacyglycerol kinase, DAG kinase; Secreted phospholipase A2, sPLA2; Autotaxin, ATX; Lipid phosphate phosphatase 1, LPP1; lecithin-cholesterol acyltransferase, LCAT). **K**. Correlation plot between aNPC proliferation and serum LPA16:0 abundance (relative values). **L**. Serum abundance of LPA16:0 in CTRL versus HR individuals (F2,56 = 3.10, p < 0.05, one-way ANOVA). **M, N**. Correlation between LPA16:0 abundance and trait (**M**) and state (**N**) anxiety levels in high-risk individuals. **O**. Serum phosphatidylcholine abundance in CTRL versus HR individuals (F2,56 = 0.99, p > 0.05, one-way ANOVA). **P**. Serum lysophosphatidylcholine abundance in CTRL versus HR individuals (F2,56 = 3.39, p = 0.04, one-way ANOVA). **Q**. Serum diacylglycerol abundance in CTRL versus HR individuals (F2,56 = 3.11, p = 0.05, one-way ANOVA). **R**. Indirect measure of ATX activity in serum from CTRL versus HR individuals (F2,56 = 3.355, p = 0.042, one-way ANOVA). **S**, Indirect measure of LPP1 activity in serum from CTRL versus HR individuals (F2,56 = 0.738, p > 0.05, one-way ANOVA). **T**. aNPC proliferation after treatment with serum from CTRL, HR-S and HR-S + KI6425 (CTRL vs. HR-Susc.; t14=3,287, p = 0.0054, unpaired t-test, two-tailed, n = 8 per group, HR-Susc+KI6425 vs. HR-Susc.; t14=2,332, p = 0.0352, unpaired t-test, two-tailed, n = 8 per group). **U**. aNPC proliferation after cLPA16:0 treatment (F4,25 = 10.17, p < 0.0001, one-way ANOVA; n = 3 wells per condition for each replicate, 2 independent replicates). For correlation plots, linear regression, r^2^ and p values are indicated in the graphs. Histograms show average ± SEM. * p<0.05; *** p<0.001. **V.** UMAP representation of two integrated human hippocampus single-cell RNA sequencing datasets^51,50^ (left panel), annotated with a label transfer algorithm trained on Tran et al., 2021^50^ (right panel). **W.** Expression level for *LPAR1* marker using a feature plot (left panel) and a dot plot representation (right panel).

To identify metabolites associated with both anxiety and aNPC proliferation, we submitted serum from the BipOff cohort to untargeted mass spectrometry-assisted metabolome analysis. Three metabolites had a significant correlation with aNPC proliferation and anxiety, with a significance threshold set to p<0.01 to both correlations: Lysophosphatydic acid 16:0, Fibrinopeptide B and Lysophosphatydilserine 18:0. Of these, we focused on Lysophosphatydic acid 16:0 (LPA 16:0) as it displayed the strongest correlation with aNPC proliferation (Fig. 2J, K). Furthermore, LPA16:0 abundance was significantly increased in HR-susceptible subjects compared to both healthy controls and HR-resilient individuals (Fig. 2L) and correlated with trait- and state-anxiety (Fig. 2M, N) in HR individuals, suggesting that LPA16:0 signaling might be involved in anxiety and aNPC proliferation. Of all the analyzed LPA forms (14:0, 16:0, 16:1, 16:3, 18:0, 18:1, 18:2, 18:3, 20:4), only LPA16:0 and LPA 20:4 were detected in the serum of the BipOff cohort. There are two pathways that synthesize extracellular LPA (Fig. 2J)^53^. The first, which produces most circulating LPA^54^, involves the membrane phosphatidylcholine (PC) cleavage into lysophosphatidylcholine (LPC) through the activation of the phospholipase A (PLA1 or 2). Subsequently, autotaxin (ATX) cleaves the head group (choline) on the LPC into LPA^55^ which can be further converted to diacylglycerol (DAG) by lipid phosphate phosphatase 1 (LPP1, Fig. 2J). While there was no difference in the total levels of PC between groups (Fig. 2O), the levels of LPC and DAG were decreased in HR-susceptible individuals compared to healthy controls (Fig. 2P, Q). These modifications in metabolites abundance resulted in increased LPA:LPC ratios (Fig. 2R) but not DAG:LPA ratios (Fig. 2S), suggesting an increased activity of the LPA-producing ATX in HR-susceptible individuals.

There are 6 known LPA receptors. LPA_1_, LPA_2,_ LPA_4_ and LPA_6_ are expressed in the hippocampus with LPA_1_ most strongly expressed in adult neural stem cells while LPA3 and LPA_5_ are undetectable^56,57^. To gain insight in LPA16:0-LPA1 signaling, we used the BBA assay and incubated aNPC with serum from either healthy controls or HR-susceptible individuals in presence of the LPA_1_ antagonist Ki16425. While serum from susceptible individuals decreased aNPC proliferation as previously observed, Ki16425 treatment normalized cell proliferation to levels observed in healthy controls (Fig. 2T), suggesting that LPA_1_ signaling was required for the effect of HR-susceptible patients’ serum on aNPC proliferation in the BBA assay. To examine whether LPA16:0 was sufficient to reduce aNPC proliferation, we added a non-hydrolysable, cyclic form of LPA16:0 (cLPA16:0) to cultured aNPC for 24h. cLPA16:0 treatment showed a dose-dependent, bimodal effect on aNPC proliferation (Fig. 2U), where low concentrations (30 nM) decreased aNPC proliferation, whereas higher concentrations (60 nM and 600 nM) had no effects on proliferation. Similarly, LPA16:0 decreased aNPC proliferation, albeit at a higher dose than cLPA16:0, likely due to its very short half-life (Supp. Fig. 3C). In contrast, the most studied form of LPA, LPA18:1 produced a dose-dependent increase in aNPC proliferation, that was blocked by the LPA /LPA antagonist diacylglycerol-phosphate (DGPP, Supp. Fig. 3D). Thus, LPA16:0-LPA_1_ signaling was both necessary and sufficient to explain the effect of serum from HR-susceptible patients on aNPC proliferation in the BBA assay.

To assess LPA_1_ expression in the human hippocampus, we interrogated two published snRNA-sequencing human databases containing hippocampal cells^58,59^. Following a careful quality control, we selected only high-quality cells for each dataset. Then, we used canonical correlation analysis (CCA) from Seurat to generate an integrated database (Fig. 2V, left panel) and observed a clear overlap between the different datasets. We then used a previously annotated dataset^59^ as reference to determine the identity of cells in the dataset of Ayhan et al.^58^, through cell type label transfer (Fig. 2V, right panel). These computational approaches revealed that cells expressing LPA_1_ correspond to non-neuronal cells, principally oligodendrocytes and microglia, and to a lower extent, mural cells, OPC and astrocytes (Fig 2W). To further assess the implication of LPA_1_ in human anxiety, we examined the genetic association between anxiety and single nucleotide polymorphisms (SNPs) on the genes involved in LPA signaling: *LCAT*, *PLA1A*, *ENPP2* (Autotaxin), *PLPP1* and *LPAR1* (LPA_1_). According to Geneatlas (a publicly available database reporting associations between multiple phenotypes and whole-genome SNPs in the UK Biobank^60^), 11, 122, 58, 184 and 188 SNPs within *LCAT, PLA1A, ENPP2, PLPP1* and *LPAR1* genes respectively, were associated with at least one of the three following anxiety phenotypes: neurotic, stress-related and somatoform disorders (F40-48), phobic anxiety disorders (F40), other anxiety disorders (F41)^60^. In the Lausanne University Hospital PsyMetab cohort (n=2479 patients), univariate analyses assessing associations between the SNPs found in Geneatlas and anxiety indicated a significant result for a SNP localized in the *LPAR1* gene (rs10817103C>T). This finding was confirmed by a multivariate linear mixed model, where patients carrying the T allele had a 40% lower risk to suffer from anxiety as compared to patients carrying the CC genotype (odds ratio: 0.61, 95% confidence interval: 0.43-0.86, p=0.005). *LPAR1* rs10817103C>T is associated with a decreased expression of LPA_1_^61^ and is localized in an enhancer active in neurons. Together, this genetic analysis supports our experimental results showing that LPA16:0-LPA_1_ signaling is involved in anxiety.

### The regulation of stress resilience by LPA16:0-LPA_1_ signaling requires adult neurogenesis

To explore the role of LPA16:0-LPA_1_ signaling in anxiety and stress response, we started by assessing LPA16:0 in mouse serum. To this aim, we used EPM and LDT to test trait anxiety in a new cohort of naïve mice and divided them into two groups based on their anxiety scores, i.e., LA and HA groups (Fig. 3A, B). We then collected blood samples and performed targeted metabolomics analysis of LPAs. HA mice displayed significantly higher serum concentrations of LPA16:0, LPA20:4, and LPA18:2 than LA mice, but not in LPA18:0 and LPA18:1 (Fig. 3C and Supp. Fig. 4A-D). Notably, there was a positive correlation between LPA16:0 concentration and individual anxiety score (Fig. 3D) and LPA16:0 concentration showed an accuracy of 80% at identifying HA from LA mice (Fig. 3E), suggesting that LPA16:0 had a similar role in mice anxiety as in humans. Next, we examined the effect of exogenous LPA16:0 on anxiety and neurogenesis. To this aim, mice were first assessed for anxiety using an EPM in order to distribute them in anxiety-matched groups. Mice were then treated with either cLPA16:0 or vehicle daily for 21 days, after which we assessed their anxiety-like behavior, sampled their serum, and processed their brains for histology (Fig. 3F). To investigate the impact of cLPA16:0 treatment on anxiety-related behavior, we conducted an open-field test.

**Fig. 3.**
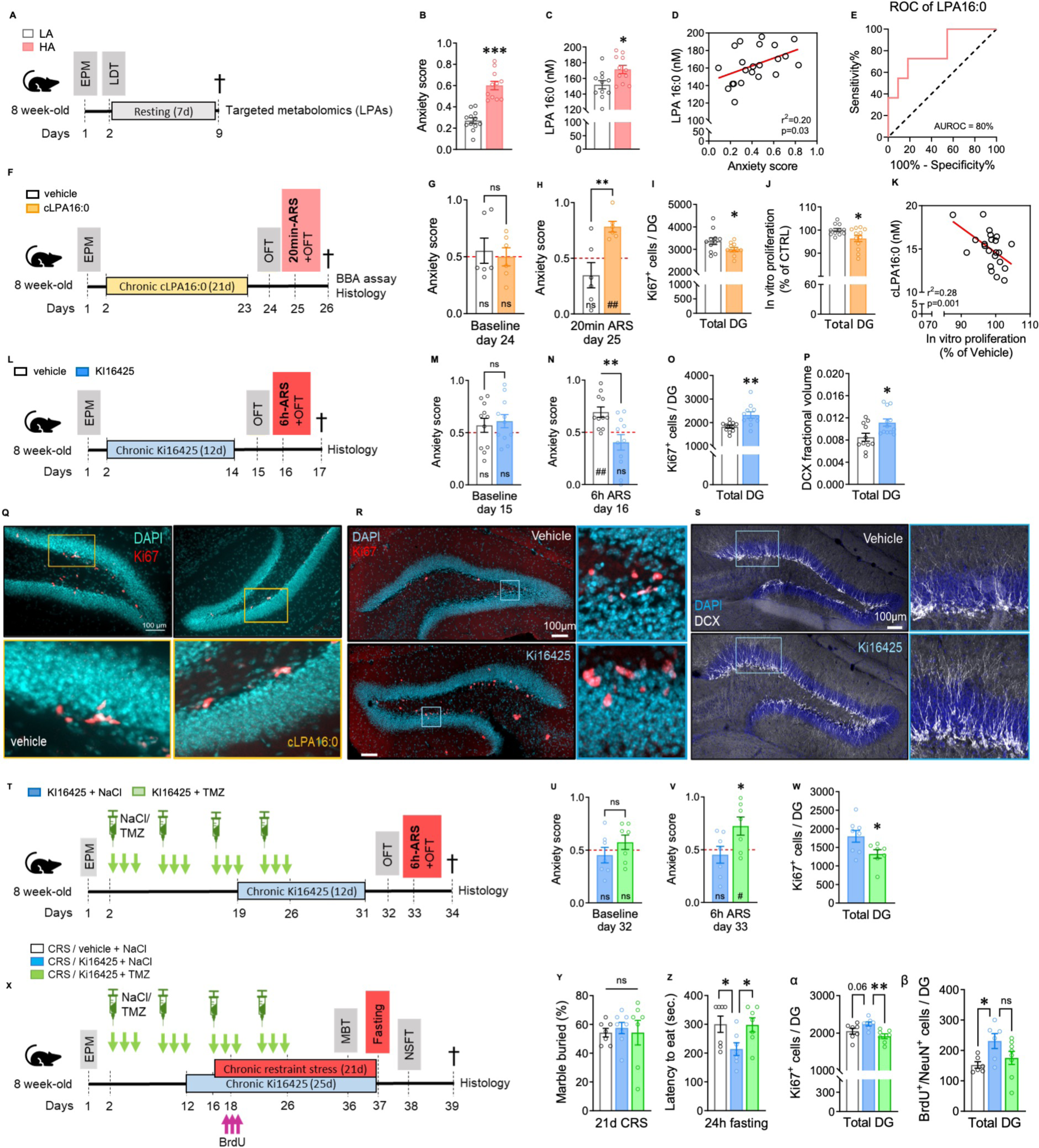
Bidirectional regulation of stress-susceptibility and neurogenesis through LPA16:0-LPA_1_ signaling. **A**. Experimental design for the anxiety trait assessment and LPA16:0 measurement. **B**. Histogram of the anxiety score of LA (white) and HA (red) mice (t21 = 7.029, p < 0.0001, unpaired t-test, two-tailed, n = 11-12 per group). **C**. Histogram of the concentration of LPA16:0 in the serum of LA vs HA mice (t20 = 2.630, p = 0.0160, unpaired t-test, two-tailed, n = 11 per group). **D**. Correlation plot between LPA16:0 serum concentrations and individual anxiety score. **E**. Receiver Operating Characteristic (ROC) curve fort the discrimination of high anxious individuals by LPA16:0. **F**. Experimental design for *in vivo* cLPA16:0 administration. **G**. Histogram of the anxiety score of mice treated with vehicle (white) and cLPA16:0 (orange) under baseline condition post treatment (t9 = 0,3932, p > 0.05, unpaired t-test, two-tailed, n =5-6 per group; vehicle: t = 0.4814, p > 0.05, One sample t test and cLPA16:0: t=0.0034, p > 0.05, One sample t test). **H.** Anxiety score following a sub-threshold 20 minutes acute restraint stress (t9 = 3.532, p < 0.01, unpaired t-test, two-tailed, n = 5-6 per group; Vehicle: t = 1.339, p > 0.05, One sample t test and cLPA16:0: t = 5.879, p < 0.01, One sample t test). **I.** Quantification of Ki-67^+^ cells in the total dentate gyrus (total DG, t20 = 2,109, p = 0.0478, unpaired t-test, two-tailed, n = 11 per group). **J.** aNPC proliferation in the BBA assay after treatment with serum from vehicle- and cLPA16:0-treated mice (t20=2.336, p = 0.0295, unpaired t-test, two-tailed, n = 11 per group). **K.** Correlation plot between serum cLPA16:0 concentration and aNPC proliferation in the BBA assay. **L.** Experimental design for *in vivo* Ki16425 administration. **M, N.** Histograms of the anxiety score of mice injected with vehicle (white) or Ki16425 (blue) after 12 days of treatment under **(M)** baseline conditions (t20 = 0.4336, p > 0.05, unpaired t-test, two-tailed; vehicle: t = 1.044, p = 0.3212, One sample t test and Ki16425: t = 1.736, p > 0.05, One sample t test, n = 11 mice per group) and **(N)** after 6 hours of ARS (t20 = 3.254, p = 0.0040, unpaired t-test, two-tailed; vehicle: t = 3.952, p = 0.0027, One sample t test and Ki16425: t = 1.288, p > 0.05, One sample t test, n = 11 mice per group). **O.** Quantification of Ki-67^+^ cells in the DG after 12 days of treatment (t20 = 3.365, p = 0.0031, unpaired t-test, two-tailed, n = 11 per group). **P.** Quantification of the proportion of pixels immunostained for DCX in the DG after 12 days of treatment (t20 = 2.721, p = 0.013, unpaired t-test, two-tailed, n = 11 per group). **Q.** Representative confocal micrographs of Ki-67 immunostaining (red) in vehicle-(left) and cLPA16:0-treated mice (right). **R**. Representative confocal micrographs of Ki-67 immunostaining (red) in vehicle-(up) and Ki16425-treated mice (down). **S.** Confocal micrographs of DCX (white) immunostaining in vehicle-(up) and Ki16425-treated mice (down). DAPI staining is shown in blue. **T.** Timeline showing the experimental design. Mice received an intraperitoneal injection of temozolomide (TMZ) or NaCl on the first 3 days of a week for 4 consecutive weeks. At the end of the third week of treatment, mice received a daily intraperitoneal injection of Ki16425. **U, V.** Histograms of the anxiety score of mice injected with Ki16425 + NaCl (blue) and Ki16425 + TMZ (green) under **(U)** baseline conditions (t12 = 1.207, p > 0.05, unpaired t-test, two-tailed; Ki16425 + NaCl: t = 0.637, p > 0.05, One sample t test and Ki16425 + TMZ: t=1.089, p > 0.05, One sample t test, n = 7 mice per group) and **(V)** after 6 hours of ARS (t12 = 2.312, p = 0.0394, unpaired t-test, two-tailed; Ki16425 + NaCl: t = 0.590, p > 0.05, One sample t test and Ki16425 + TMZ: t = 2.606, p = 0.0404, one sample t test, n = 7 mice per group). **W.** Quantification of Ki-67^+^ cells in the DG (t12 = 2,356, p = 0.0349, unpaired t-test, two-tailed, n = 7 per group). **X.** Timeline showing the experimental design. Mice received an intraperitoneal injection of TMZ or NaCl on the first 3 days of a week for 4 consecutive weeks. Ten days after the beginning of TMZ treatment, mice received a daily intraperitoneal injection of Ki16425 for 25 days. Sixteen days after the beginning of TMZ treatment, mice underwent a CRS protocol for 21 days. After 2 days of CRS regimen, mice received 3 injections of BrdU every 2 hours. **Y.** Assessment of anxiety-like behavior in a marble burying test (MBT). Histogram of the percentage of marble buried after 20minute-session of MBT (F2,18 = 1.038, p > 0.05, one-way ANOVA, n = 7 mice per group). **Z.** Assessment of depressive-like behavior in a novelty suppress feeding test (NSFT). Histogram of the latency to the first bite after 6 minutes of NSFT (F2,18 = 3.846, p = 0.0407, One-way ANOVA; *p < 0.05, Tukey’s multiple comparisons, n = 7 mice per group). **α.** Quantification of Ki-67^+^ cells in the DG (F2,18 = 6,169, p = 0.0091, One-way ANOVA; p = 0.06, **p < 0.01, Tukey’s multiple comparisons test, n = 7 mice per group). **β.** Quantification of BrdU^+^/NeuN^+^ cells in the DG (DG, F2,18 = 3,514, p = 0.05, One-way ANOVA; *p < 0.05, p^ns^ = 0.160, Tukey’s multiple comparisons test, n = 7 mice per group). Linear regression, r^2^ and p values are indicated in the graphs when significant correlations were found. Histograms show average ± SEM. * p<0.05; ** p<0.01; *** p<0.001; ns: not significant. Comparison between the group mean and the hypothetical value of 0.5, to assess anxiety withing each group are shown within each histogram bar. One-sample t-test. #: p<0.05; ##: p<0.01.

Surprisingly, mice treated with cLPA16:0 did not exhibit any significant changes in baseline anxiety-like behavior compared to the vehicle-treated mice (Fig. 3G). Considering our previous findings in the human cohort, where LPA16:0 levels were elevated only in individuals at high risk and susceptible to anxiety disorders (Fig. 2L), we postulated that LPA16:0 might trigger an exaggerated stress response. To test this hypothesis, we subjected both cLPA16:0- and vehicle-treated mice to a subthreshold, 20 minutes acute restraint stress (ARS), a paradigm known to unveil a susceptible phenotype by lowering the animals’ stress reactivity threshold^62,63^. Mice treated with cLPA16:0 displayed increased anxiety-like behavior following 20 min. ARS compared to the control group (Fig. 3H), indicating that cLPA16:0 treatment exacerbated stress reactivity. Furthermore, cell proliferation in the dentate gyrus, assessed by immunohistochemistry against the endogenous cell proliferation marker Ki-67, was reduced in mice treated with cLPA16:0 (Fig. 3I, Q). Similarly, serum from cLPA16:0-treated mice decreased aNPC proliferation in the BBA assay as compared to serum from vehicle-treated mice (Fig. 3J) and serum cLPA16:0 concentration negatively correlated with aNPC proliferation in the BBA assay (Fig. 3K). Thus, increased circulating LPA16:0 impaired hippocampal neural stem/progenitor cell proliferation and reduced stress resilience.

We then tested whether antagonizing the LPA_1_ receptor might confer stress resilience. To this aim, we tested the resilience of mice to 6 hours of ARS, which is a protocol that induces anxiety-like behavior in naïve mice. Anxiety-matched mice were injected daily for 12 days with either Ki16425 (i.p. 5 mg/Kg body weight) or vehicle and then assessed for their basal anxiety using an open-field test. One day later, they were subjected to 6 hours of ARS followed by an open-field test (OFT) 20 minutes afterwards to assess state anxiety (Fig. 3L). The anxiety score before ARS did not differ between Ki16425- or vehicle-treated mice (Fig. 3M). After ARS however, vehicle-treated mice showed an expected increase in anxiety score, whereas mice treated with Ki16425 exhibited no anxiety-like behavior (Fig. 3N). The resilience profile exhibited by the Ki16425-treated mice was associated with an increase in aNPC proliferation (Fig. 3O, R) and an increase in the expression of the immature neuron marker DCX in the dentate gyrus (Fig. 3P, S). To test the reliability and reproducibility of our findings, we conducted a replication experiment in a different laboratory, with a different experimenter. Consistently, in this separate experiment, Ki16425 administration resulted in a significant enhancement of stress resilience from a 6hour-ARS paradigm with no change in anxiety under baseline conditions (Supp. Figure 4E-G). Thus, antagonizing LPA_1_ increased adult neurogenesis and resilience to an acute stress.

To test whether adult neurogenesis may mediate the stress resilience effects of Ki16425, we used the anti-mitotic drug temozolomide (TMZ) to suppress adult neurogenesis. Anxiety-matched mice were first treated with either NaCl or TMZ for 4 weeks, 3 days/week followed by, after the third week of TMZ treatment, 12 days administration of Ki16425 and then assessed for resilience to a 6h-ARS (Fig. 3T). Before ARS, we found no group difference in anxiety in an open field test (Fig. 3U). After ARS, Ki16425+NaCl-treated mice did not display increased anxiety, confirming the resilience profile observed previously (Fig. 3N). Strikingly, mice treated with both Ki16425 and TMZ showed increased anxiety (Fig. 3V), indicating a decreased stress resilience in the TMZ-treated group. In addition, Ki16425+TMZ-treated mice showed less cell proliferation than Ki16425+NaCl-treated mice (Fig. 3W), confirming the effect of TMZ on the reduction of adult neurogenesis. These results suggest that the effect of Ki16425 on acute stress resilience required intact adult neurogenesis.

The experiments described above tested the effect of LPA_1_ antagonism on anxiety after an acute stress. To test the effect of Ki16425 on depressive-like behavior after a chronic stress, we repeated these experiments with 21 days of CRS. To this aim, anxiety-matched mice were treated for 4 weeks with either NaCl or TMZ. Starting after the second cycle of TMZ (11 days after the first day of TMZ treatment), mice were injected daily with Ki16425 for 25 days and, starting 4 days after treatment initiation, were all subjected to CRS for 21 days. Two days later, mice were injected with BrdU. One day before the end of the CRS, anxiety-like behavior was assessed on a marble burying test (MBT) and, one day later, depressive-like behavior was tested on a novelty-suppressed feeding test (NSFT) (Fig. 3X). We found no difference between groups in the number of marbles buried, suggesting no difference in anxiety (Fig. 3Y). However, in a NSFT, mice treated with Ki16425 showed a decreased latency to feed as compared to vehicle-treated animals, an effect that was abolished by TMZ treatment (Fig. 3Z). Following 21 days of CRS, Ki16425 increased cell proliferation (Fig. 3α) and the number of newly formed neurons (Fig. 3β), as compared to NaCl, an effect that was abolished by TMZ. Together, these results indicate that the increase in adult neurogenesis was required for the antidepressant effect of Ki16425 after stress.

### Platelet depletion reduces circulating LPA16:0 and increases stress resilience and adult neurogenesis

Serum LPA is mostly produced by platelets^64–67^. Consistent with this primary source, thrombocytopenia is characterized by a significant reduction in circulating LPA^67,68^. We therefore assessed whether LPA16:0 may be reduced by decreasing platelets. To this aim, we injected 3 mice with anti-platelet or control serum for 2 days and analyzed their plasma by pooling all animals from each condition. Anti-platelet treatment reduced platelets and LPA16:0 to undetectable levels (Fig. 4A). We then tested the possibility that anti-platelets serum may regulate adult neurogenesis and stress resilience. To this aim, anxiety-matched mice were administered with anti-platelet serum every 2 days for 20 days^69,70^ and tested for anxiety in an OFT, followed by a stress-resilience test using a 6h ARS (Fig. 4B). Platelet-depleted mice exhibited reduced anxiety-like behavior under baseline conditions whereas mice injected with control serum displayed a normal anxiety score (Fig. 4C). After ARS, control mice showed an expected increase in anxiety score, whereas mice treated with antiplatelets exhibited no anxiety-like behavior (Fig. 4D). This resilience profile of platelet-depleted mice was associated with an increase in the proliferation of aNPCs in the DG (Fig. 4E). Thus, depleting platelets eliminated circulating LPA16:0 and mimicked the stress-resilience effect of LPA1 antagonism.

**Fig. 4.**
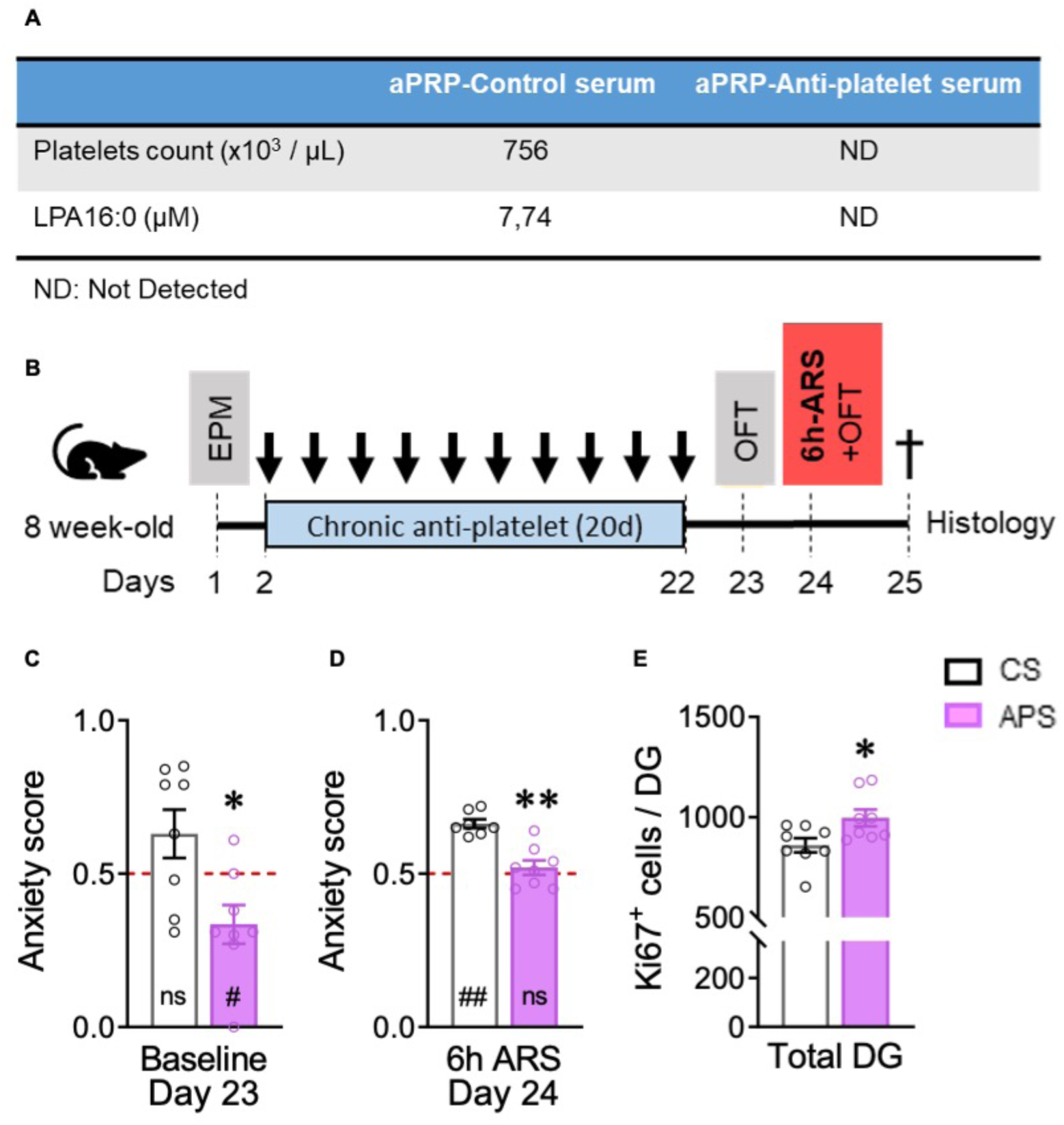
Platelet depletion reduces circulating LPA16:0 and increases stress resilience and adult neurogenesis. **A.** Table showing platelets count and levels of LPA16:0 in activated platelet-rich plasma from mice treated with an anti-platelet serum or control serum for 2 days. **B.** Timeline showing the experimental design. Mice received an intraperitoneal injection of anti-platelet serum or control serum every other day for 20 days, followed by assessment of resilience to a 6 hours ARS. **C, D.** Histograms of the anxiety score of mice injected with control (white) or anti-platelet serum (purple) after 20 days of treatment under **(C)** baseline conditions (t14 = 2.922, p = 0.0011, unpaired t-test, two-tailed; control: t = 1.651, p > 0.05, One sample t test and anti-platelet: t = 2.610, p = 0.0349, One sample t test, n = 8 mice per group) and **(D)** after 6 hours of ARS (t14 = 5.005, p = 0.0002, unpaired t-test, two-tailed; control: t = 11.29, p < 0.0001, One sample t test and anti-platelet: t = 0.852, p > 0.05, One sample t test, n = 8 mice per group). **E.** Quantification of Ki-67^+^ cells in the DG (t14 = 2,452, p = 0.0279, unpaired t-test, two-tailed, n = 8 per group). Histograms show average ± SEM. * p<0.05; ** p<0.01; *** p<0.001; ns: not significant. Comparison between the group mean and the hypothetical value of 0.5, to assess anxiety withing each group are shown within each histogram bar. One-sample t-test. #: p<0.05; ##: p<0.01

## Discussion

In the present study, we found that spontaneously anxious mice exhibited reduced cell proliferation and adult neurogenesis in the dentate gyrus of the hippocampus. Using an *in vitro* model of adult neurogenesis, the BBA assay, we observed that serum from spontaneously anxious mice, experimentally stressed mice or human patients suffering from anxiety, reduced aNPC proliferation proportionally to their anxiety levels. Untargeted mass spectrometry identified that the serum concentration of LPA16:0, a signaling lipid, correlated with anxiety in humans and mice as well as with the effect of their serum on aNPC proliferation. LPA16:0 was necessary and sufficient to replicate the effect of human serum on the BBA assay, which was mediated by the LPA_1_ receptor. In a combined analysis of the Geneatlas and an in-house cohort of patients, we found an association between anxiety and the rs10817103C>T SNP on the enhancer region of the *LPAR1* gene, which is known to decrease LPA_1_ expression. Furthermore, LPA16:0-LPA_1_ signaling in mice bidirectionally regulated stress resilience and adult neurogenesis and the increase in stress resilience produced by LPA_1_ antagonism was mediated by adult neurogenesis. Finally, reducing platelets, the main source of LPA in blood, reduced LPA16:0 and induced stress resilience along with increasing aNPC proliferation in the dentate gyrus. Together, these results highlight a crucial role of platelets in mediating stress susceptibility in anxious individuals, by the production of LPA16:0 and the inhibition of adult hippocampal neurogenesis.

Growing evidence indicate that blood-circulating molecules regulate brain plasticity, as demonstrated by blood transfusions or parabiosis experiments ^33–38^. To circumvent the technical difficulties of these experiments, we have developed a simple *in vitro* model of adult hippocampal neurogenesis, named the BBA assay, that enables the discovery of blood-circulating molecules that regulate adult hippocampal neurogenesis in mice. The BBA assay exploits the sensitivity of aNPC to molecular changes in the neurogenic niche that is enabled by the extended repertoire of membrane receptors they express^71^. Due to the difficulty of assessing adult neurogenesis in the human brain^72–77^, the relevance of the BBA assay for human adult hippocampal neurogenesis was not addressed here. However, the human hippocampus expresses LPA_1_ mRNA in numerous cell types including oligodendrocytes, microglia, astrocytes and mural cells (Fig. 2W). The shared repertoire of receptors between aNPC and niche cells indicates that the BBA assay may be used as a cell-based biomarker of the integrity of the hippocampal neurogenic niche, which is relevant to aspects of hippocampal function such as emotional regulation. The present study highlights the translational value of the BBA which, combined with untargeted metabolomics and clinical assessment, provides an entry point for the identification of novel mechanisms of regulation of adult neurogenesis by circulating molecules in the context of anxiety disorders. Similar approaches have been used to study the role of circulating molecules in other diseases related to hippocampal dysfunction such as Alzheimer’s disease^78,79^ or depression^80^.

The different forms of LPA play a role in several cellular processes that include smooth muscle contraction^81^, platelet aggregation^82^, cell growth and differentiation^83^ and emotional regulation^84^. These effects are principally mediated by 6 G protein-coupled receptors which are differentially expressed in various cell types and display differential affinities for the forms of LPA^53,55^. Previous human studies have shown an association between mood disorders and LPA signaling: Genome-wide microarray studies have found that the expression of a panel of genes that included LPA_1_ is associated with mood disorders^85,86^. Lipidomic analysis of serum from major depressive disorder patients found a decreased abundance of LPA22.6^87^ and a more recent study found that changes in several triglycerides and LPA16:1 are sufficient to distinguish currently depressed patients from remitted patients or healthy controls^88^. Similarly, the levels of autotaxin in the serum and cerebrospinal fluid were reduced in patients affected with major depressive disorder and serum autotaxin was restored upon electroconvulsive treatment^89^. These reports are consistent with our observation of an association between anxiety and circulating LPA16:0 as well as with a SNP on LPA_1_ and support the view that LPA-LPA_1_ signaling plays a role in anxiety and stress susceptibility in several psychiatric diseases.

Several mouse studies explored the role of LPA-LPA_1_ signaling in brain function and mood disorders and found, in contrast with our observations, that LPA_1_ activation increases adult neurogenesis and produces antidepressant effects: First, the spontaneous LPA_1_ null mice display depression-like behavior and decreased adult hippocampal neurogenesis^90,91^. Yet, LPA_1_ deficient mice suffer from perinatal mortality and are subject to compensatory adaptations that may interfere with the expression of other LPA receptor isoforms. It is therefore unclear whether the phenotype of the LPA_1_ null mice exclusively results from impaired LPA signaling^92^. Second, the pharmacological blockade of LPA_1_ by Ki16425 was found to increase anxiety- and depression-like phenotypes^93–95^ and inhibit adult neurogenesis^95^. However, differences in the mode of administration between these reports and the present study (intracerebroventricular and intraperitoneal, respectively), likely underlie the differences in biological effect. Finally, the intracerebroventricular administration of LPA18:1 was reported to increase adult neurogenesis^95,96^ and to reduce anxiety- and depression-like behavior^93,95,96^ (but see^97^), as well as autotaxin overexpression in the hippocampus^98^. However, LPA18:1 produces opposite effects of LPA16:0 on aNPC proliferation (Fig. 2U, Supp. Fig. 3C, D), likely resulting from differences in affinities to the LPA_1_ receptor. Thus, the net effect of LPA on adult neurogenesis and stress resilience likely depends on the relative abundance of the different forms of LPA in the periphery and in the central nervous system. However, in the present study, the selection of serum metabolites based on the double correlation (with cell proliferation in the BBA assay and trait anxiety in patients), highlighted the role of circulating LPA16:0 in anxiety disorders. This suggests that mechanisms involved in the production of circulating LPA16:0 may regulate adult neurogenesis.

Since they constitute the main source of LPA in the blood^64–68^, platelets are expected to play a major role in the regulation of adult neurogenesis and stress resilience. Furthermore, LPA16:0 was found to be among the most strongly upregulated lipids in activated human platelets^65^, supporting our observations that platelets-deprived mice display undetectable levels of LPA16:0 (Fig. 4A). Consistently, earlier studies described an association between psychiatric diseases, including major depression, and increased platelets volume and platelet reactivity^100,101^, suggesting that platelets play an active role in mood disorders.

Together, the experimental evidence presented here suggests that platelets in anxious individuals produce circulating LPA16:0 that target the LPA_1_ receptor, leading to reduced adult neurogenesis and impaired stress resilience. This mechanism may contribute to the pathophysiology of many psychiatric conditions and offer novel therapeutic avenues. In addition, the BBA assay presented here, together with LPA16:0 measurements may provide a diagnostic tool to assess anxiety in psychiatric patients, paving the way for personalized treatment.

## Materials and Methods

### The blood-brain axis (BBA) assay

*Cell culture:* aNPC were isolated from the dentate gyrus (DG) of adult Fisher 344 rats and cultured in medium Dulbecco’s Modified Eagle Medium (DMEM/F12) supplemented with N2 and FGF-2 (20ng/ml) as previously described^102^. They were plated at a density of 60,000 cells per well (total volume = 500µl) in a 24-well-plate coated with Poly-D-lysine hydrobromide at 0.1mg/ml (P6407 sigma-aldrich) and Laminin at 20µg/ml (23017015 Life technologies). Twenty-four hours after plating, the medium was removed and replaced with fresh medium containing 0.2% serum for 24h. The medium was then supplemented with 5 μM BrdU for 30 min and washed twice with pre-heated fresh medium and fixed with 4% paraformaldehyde for 30 min. The plate was briefly washed with PBS 0.1M and stored at 4°C until used. Immunocytochemistry against BrdU and/or Ki-67 was processed as follows: BrdU immunohistochemistry was preceded by DNA denaturation with 2 M HCl for 15 min at 37°C, and rinsed in 0.1 M borate buffer pH 8.5 for 15 min. Then, aNPC were incubated in blocking solution containing 0.3% Triton-X100 and 10% deactivated horse serum (Gibco, 16050122) for one hour. Plates were incubated at 4°C with mouse monoclonal anti-BrdU (24 h, 1:250, Abcam) or Rabbit anti-Ki-67 (1:300, Abcam, ab15580) followed by goat anti-mouse 488 (1:300, Invitrogen) or goat anti-rabbit 488 (1:300, Invitrogen), respectively for 1 h at room temperature. After immunostaining, 10 minutes incubation into 4,6 diamidino-2-phenylindole (DAPI, 1:500) was used to reveal nuclei.

For neurosphere cultures, aNPC were grown with 0.2% serum at a density 60,000 cells per well in a 24-well-plate under floating conditions and were incubated at 37°C for 3 days. The neurospheres were counted and their size measured using an inverted light microscope and the ImageJ software. Control cultures were treated with serum-free medium.

*Image acquisition and analysis for in vitro experiments:* Images were acquired using an Eclipse Ti2 inverted microscope (Nikon). The number of BrdU-labeled aNPC was counted in one randomly selected large field (tiles 4x4) in each well of the plate. For the validation of the BBA assay (Fig. 1B), we used 2-3 wells per condition in 3 independent replicates. For all other experiments, each well stands for an experimental unit and was replicated at least twice using 2 independent cohorts of mice for each condition (trait and state conditions). The number of BrdU-labeled aNPC was compared with the total number of aNPC in each selected field to obtain a proliferation ratio, which was then normalized to the control group values to enable comparisons across conditions.

*Cell viability:* Cell viability was evaluated using the WST-1 assay. Cells (6.0 × 10^4^ cells/well) were grown in 24-well plate. After exposure to serum for 24h., 25 μl cell proliferation reagent WST-1 (Roche, Cat. No. 05015944001, Switzerland) was added to each 500 μL/well serum-free DMEM/F12 (0.5:10 dilution), followed by incubation for 2 hours at 37°C. Absorbance was measured at 440 nm using a microplate reader (Tecan, Männedorf, Switzerland).

### Mice

All experiments were performed with the approval of the Cantonal Veterinary Authorities (Vaud, Switzerland) and carried out in accordance with the European Communities Council Directive of 24 November 1986 (86/609EEC). All experiments were performed on C57Bl/6J mice obtained from Charles River Laboratories. After arrival, animals were housed four per cage and allowed to acclimate to the vivarium for one week. All animals were subsequently handled for 1 min. per day for a minimum of 3 days. Animals were weighted upon arrival as well as weekly to monitor health. Mice were maintained under standard housing conditions on corncob litter in a temperature-(23 ± 1°C) and humidity-(40%) controlled animal room with a 12h. light/dark cycle (8–20 h.), with unlimited access to food and water.

### Chronic and acute restraint stress

This protocol involved 21 days of chronic retrain stress (CRS) as previously described^44,103^. Animals were randomly assigned to either the control or CRS group. Animals were introduced headfirst into 50 ml Falcon tubes (11.5 cm in length; diameter of 3 cm) from which the cap was removed, and the bottom was perforated with four 0.4 cm holes to enable breathing. Tissue was added at the caudal extremity to adjust the physical constraint to the mouse size and to allow the tail to expand out of the tube. The mice were subjected to this restrained environment for two consecutive hours every day for a period of 21 days. Control mice were left undisturbed in their home cage except for handling and body weighting each day for 21 days. The acute restraint stress protocol is based on the protocol as previously described^63^. Animals of the stress group were restrained once, using the protocol described for the chronic stress. After one 20-min-restraining or 6-hours-restraining period, mice were transferred into their home-cage for an additional 20 min interval followed by an open-field test to assess anxiety-like behavior.

### Behavioral tests

*Open-field test (OFT):* The test was performed as previously described^103,104^. The apparatus consisted of a square Plexiglas arena (40 x 40 x 40 cm) that was illuminated with dimmed lights (25 lx). The floor was cleaned between each trial to eliminate olfactory cues. Mice were introduced facing the wall of the arena and allowed to freely explore the arena for 10 min. A virtual center zone (15 x 15 cm), thigmotaxis zone (30 x 30 cm) and an intermediate zone were included for the behavioral analysis as indicator for anxiety-like behavior. A video tracking system (Anymaze) recorded the path of each mouse as well as the total distance travelled, and the time spent exploring each zone.

*Elevated plus maze test (EPM):* The test was performed as previously described^104^. The apparatus was made from black PVC with a white floor. The apparatus consisted of a central platform (5 x 5 cm) elevated from the ground (65 cm) with two opposing open (30 x 5 cm) and two opposing (30 x 5 x 14 cm) closed arms. Light conditions were maintained at 14–15 lux in the open arms, and 3–4 lux in the closed arms. Animals were placed at the end of the closed arms facing the wall, after which the animals were allowed to freely explore the apparatus for 5 min. Mice were tracked (Anymaze) to measure the time spent in each arm and in the risk zones (edge of the open arms).

*Light/Dark Test (LDT):* The LDT was performed as previously described^40^. A 60 x 40 x 21 cm high Plexiglas box was divided into a dark (20 x 40 x 21 cm) and a light (40 x 40 x 21 cm; 400 lux) compartments separated by an open door (5 x 5 cm) located in the center of the partition at floor level. Each mouse was placed into the light chamber facing the door. Mice were allowed to freely explore the apparatus for 6 minutes. The Anymaze software was used to analyze anxiety-like behavior by calculating the time spent in each zone.

*Marble burying test (MBT):* As previously described^105^, the apparatus consists of an open transparent plastic box (40 x 25 x 20 cm) filled around 6 cm deep with bedding material across the whole cage. Twenty dark marbles (diameter: 16 mm) are spaced evenly in a 4 x 5 grid on the bedding. Mice were given 20 minutes to freely explore the cage (400 lux). At the end, the mice are removed, and every buried marble (more than 2/3 covered by litter) is counted. The number of buried marbles is used as an indicator of anxiety.

*Novelty suppressed feeding test (NSFT):* As previously described^106^. The apparatus consists of a square Plexiglas arena (40 x 40 x 40 cm; > 400 lx). The floor was layered with approximately 2 cm of wood bedding. In the center of the square, a single pellet of food was placed on a white paper circular taped to a tissue culture disk (20 x 100 mm). Twenty-four hours before the test, all the food was removed from the home cage to food restricted the mice. The mice were introduced into the arena, facing the wall, and given 3 minutes for free exploration. Anymaze video tracking system recorded the movement trajectory of each mouse, and the videos were analyzed to assess the latency for the first bite. After the test, the food pellet was removed, and the mice were then returned to their cage with food and the amount of food consumed in 20 minutes was measured (in-cage consumption).

*Anxiety scores:* The anxiety score was calculated as previously described^104,107^ with the average of standardized scores of each anxiety-related behavior tests. For Fig. 1H, 1Q and 3B, this score included the relative time spent in the dark chamber of the LDT and the relative time spent in the closed arm of the EPM. For Fig. 3F-U and figure 4C, D, this score included the relative time spent in thigmotaxis and in the center of the arena of an open-field. Standardization consisted in subtracting the minimum value of the whole population to the value of each animal and dividing the result by the maximum value of the whole population minus the minimum value of the whole population: (x – min value) / (max value – min value). This procedure yields scores which are distributed along a scale from 0 to 1, where 1 reflects high anxiety. To assess anxiety levels in each group, a one-sample t-test was employed, using 0.5 as the hypothetical threshold value as scores were distributed along a scale from 0 to 1. If the mean anxiety score of a group exceeded 0.5, it was classified as anxious; if the score was below 0.5, the group was considered non-anxious. This approach allowed for a statistical comparison of each group’s anxiety levels against the predefined baseline of 0.5, facilitating clear distinctions between anxious and non-anxious subjects.

### Mouse blood collection and preparation

Blood was collected (Multivette® 600 µl, Clotting Activator/Serum) by intracardiac punction using a 1ml syringe with a 21G7/8 needle after pentobarbital (10 ml/kg, Sigma-Aldrich, Buchs, Switzerland) anesthesia before mouse perfusion. After sampling, the blood was left undisturbed at room temperature for 15 minutes to enable clotting. The clot was removed by centrifuging at 1,500 x g for 10 minutes in a refrigerated centrifuge. The resulting supernatant (i.e., serum) was transferred into a clean polypropylene tube using a Pasteur pipette. The samples were apportioned into 0.5 ml aliquots and stored at - 80°C.

### Preparation of platelet-rich plasma (PRP) after anti-platelet serum injection

Mice were injected with an antiplatelet serum or a control serum for 2 consecutive days. After pentobarbital (10 ml/kg, Sigma-Aldrich, Buchs, Switzerland) anesthesia, whole blood was collected transcardially using a 21G7/8 needle containing 10% of sodium citrate, then transferred into plastic tubes. The platelet count was measured using 20 µl of whole blood in a Sysmex XE-2100™ automated hematology system. Blood samples were centrifuged at 1900 g for 1 minute with no brake at room temperature. The plasma and buffy coat were collected as PRP into a new tube. PRP was stored at -20°C for 15 min to activate platelets. Samples were then used for targeted LPA16:0 quantifications. The platelet counts and LPA16:0 levels were determined from a pool of 4 mice per condition, i.e., control serum- and antiplatelet serum-treated groups.

### Corticosterone measurement

Serum (6µL) was prepared according to manufacturer’s instructions to measure free corticosterone concentrations using an ELISA kit (Enzo Life Sciences, ADI-901-097). The method plots the standards versus hormone concentrations using linear (y) and log (x) axes and performs a 4-parameter logistic fit. Concentration of samples were then calculated from the fit.

### BrdU administration

Mice were injected intraperitoneally with 5-bromo-2-deoxyuridine (Sigma-Aldrich, Buchs, Switzerland) at a concentration of 100 mg/kg in saline (final dilution at 10 mg/ml), 3 times per day at 2-h. intervals in a single day. For proliferation studies, mice were sacrificed 24 h. after the last BrdU injection.

### Tissue collection and preparation

Mice received a lethal dose of pentobarbital (10 ml/kg, Sigma-Aldrich, Buchs, Switzerland) and were perfusion-fixed with 50 ml of 0.9% saline followed by 50 ml of 4% paraformaldehyde (Sigma-Aldrich, Switzerland) dissolved in phosphate buffer saline (PBS 0.1 M, pH 7.4). Brains were then collected, post-fixed overnight at 4 °C, cryoprotected 72h in 30% sucrose and slowly frozen on dry ice. Coronal frozen sections of a thickness of 40 μm. were cut with a microtome-cryostat (Leica MC 3050S) and slices were kept in cryoprotectant (30% ethylene glycol and 25% glycerin in 1× PBS) at −20 °C until being processed for immunohistochemistry.

### Brain slices immunohistochemistry

Immunochemistry was performed as previously described^108^. For BrdU/NeuN immunostaining, sections were washed 3 times in PBS 0.1M. BrdU detection required formic acid pre-treatment (formamide 50% in 2× SSC buffer; 2× SSC is 0.3M NaCl and 0.03 M sodium citrate, pH 7.0) at 65 °C for 2 h followed by DNA denaturation for 30 min in 2 M HCl for 30 min at 37°C and rinsed in 0.1 M borate buffer pH 8.5 for 10 min. Then, slices were incubated in blocking solution containing 0.3% Triton-X100 and 10% deactivated horse serum (Gibco, 16050122) for 1 h. After blocking, samples were incubated with primary antibodies, at 4°C overnight and then incubated with secondary antibodies for 1 h. For Ki67 and DCX immunostaining, slices were incubated in blocking solution (0.3% TritonX-100 and 10% horse serum in PBS) at room temperature for 1 h and then incubated under agitation at 4° C overnight in a blocking solution containing primary antibodies. After being washed again in PBS, sections were incubated for 2 hours in secondary antibodies. After immunostaining, slices were incubated for 10 min into DAPI (1:500) to reveal nuclei. Sections were mounted onto glass slides and cover-slipped using FluorSave (Millipore). One out of 6 sections (for a total of 10 sections) were imaged using a Nikon NI-E spinning disk fluorescence microscope. For BrdU analyses (Supp. Fig. 2A), a standardized value was calculated as previously described using the formula (x – min value) / (max value – min value). This procedure yields scores which were distributed along a scale from 0 to 1 for each parameter, 1 reflecting high *in vivo* proliferation, which were then averaged and used to evaluate *in vivo* proliferation. Individual scores were then normalized to the low anxiety (LA) group values. For DCX analysis, we assessed the proportion of immunostained pixels on each image. Data is expressed either as pixels/area (i.e. region of interest, Supp. Fig. 2), or fractional volume (Fig. 3P). For the pixels/area analysis, images were acquired using a digital camera (3CCD Hitachi HV-F202SCL) on a slide scanner microscope (×20 objective, Zeiss axioscan Z1). The number of DCX-immunostained pixels was quantified in similar regions of interest, in arbitrary units as the mean of all isolated pixels of soma. Each optical density was normalized by the subtraction of a slide section in which signal was absent (black) using image analysis software Zen 2 (black 8.0 edition and blue 2012 edition). The area of the DG was also measured using Zen blue edition software and no change in DG area was found between groups (data not shown). For the fractional volume analysis of DCX staining (Fig. 3P), images were acquired using a Nikon NI-E spinning disk fluorescence microscope of similar regions of interest between slices. We calculated the fraction of the total volume that is occupied by a DCX-positive signal in each section. The mean fractional volume per animal was derived from the average values of all sections analyzed. For Ki67 immunostaining analyses, the total number of immunostained cells in the DG was counted and multiplied by 6 to obtain total values/DG.

### Reagents and antibodies

For primary antibodies: mouse monoclonal anti-BrdU (1:250, Chemicon International, Dietikon, Switzerland), rabbit anti-DCX (1:500, Cells Signaling Technology, 4604S), rabbit anti-NeuN (1:1000, Abcam, EPR12763), rabbit anti-Ki-67 (1:300, Abcam, ab15580); rabbit anti-cleaved caspase3 (1:500, cell signaling 9579). For secondary antibodies: goat anti-mouse Alexa-594 (1:300, Invitrogen), goat anti-rabbit 594 (1:300, Invitrogen). For drugs: staurosporine (STS, from Cell Signaling Technology, 9953S), cyclic LPA16:0 (0.01 mg/ml in PBS 0.1M and 0.1% BSA fatty acid free, Polar Avanti, 7999268-68-1), DGPP (50µM in ddH_2_0 and 5% ethanol, Polar Avanti, 474943-13-0), Ki16425 (0.5 mg/ml in ddH_2_0 and 10% ethanol, Cayman Chemical, 10012659), temozolomide (TMZ, from Sigma-Aldrich, T2577, (25 mg/kg; 2.5 mg/ml in 0.9% NaCl i.p.), and rabbit anti-platelet serum (1:2 in PBS, 200 µl per mouse, Accurate Chemical & Scientific Corporation).

### Human participants

*Participants from the BipOff cohort:* The research was conducted according to the principles of the Declaration of Helsinki and approved by the University of Geneva research ethics committee (CER 13-081). All participants gave written informed consent prior to assessment. Offspring of patients were recruited after their parents gave a formal consent to contact their children. We obtained blood samples from 66 individuals from the BipOff cohort of the Geneva University Hospital: 35 offspring at high-risk (HR) of developing a psychiatric disorder, defined as having at least one parent with either ADHD or Borderline Personality Disorder (BPD) or Bipolar Disorder (BD) and 31 healthy control subjects with no history of neuropsychiatric disorder among their parents (CTRL). The offspring were between 15 and 25 years of age at inclusion, and controls between 15 and 30 years old. Control subjects were matched for age, gender, laterality and years of education. Parents were outpatients from the specialized program for BPD/ADHD/BD patients at Geneva University Hospital, where structured diagnostic assessment was conducted. Psychiatric diagnoses in all subjects were established by trained psychologists using the Diagnostic Interview for Genetic Studies (DIGS) or the Kiddie Schedule for Affective Disorders and Schizophrenia (K-SADS) under 18 years of age. This psychiatric assessment led to the stratification of high-risk individuals into two groups; HR-susceptible (n=21) or HR-resilient (n=14), based on whether they had developed a neuropsychiatric disorder at the time of evaluation. Inclusion criteria were based on age, history of psychiatric or neurological treatment for the subjects and their parents, as assessed during the interview of the subjects. Exclusion criteria for all participants were a history of head injury, current alcohol or drug abuse, daily medication. Assessment of anxiety was conducted through a classical self-report questionnaire, the State-Trait-Anxiety Inventory (STAI)^109^. The STAI is a validated 40-item self-report questionnaire that measures both state (20 items) and trait (20 items) general anxiety. Trait anxiety scores were assessed during the evaluation visit 1-3 weeks prior to lab visit. State anxiety scores were assessed immediately prior to blood sampling. Blood samples (5ml) were kept on ice for about 1 h and were left for clotting for 30 minutes, followed by centrifugation (5 min at 2300g at room temperature). Samples were then aliquoted and frozen at -20°C for 1-4 hours and then stored at -80°C.

*Participants from the PsyMetab cohort:* Since 2007, a longitudinal observational pharmacogenetic study (referred to as PsyMetab) approved by the Swiss Association of Research Ethics Committees is ongoing in the Department of Psychiatry of the Lausanne University Hospital. Patients starting a psychotropic treatment associated with a risk of developing metabolic disturbances (i.e., antipsychotics, mood stabilizers or some antidepressants)^110^ were included. Clinical data including concomitant medication prescriptions were collected during hospitalization or in outpatient centers during medical examinations based on the department guideline for the metabolic follow-up of psychotropic drugs^111^. In the present study, drug prescription was used to assess anxiety. Patients who were prescribed any benzodiazepine specifically used for treating anxiety symptoms (i.e. alprazolam, bromazepam, clorazepate, ketazolam, lorazepam, oxazepam, prazepam or chlordiazepoxide, as well as clobazam or diazepam if not co-prescribed with any antiepileptic agent) were considered as suffering from anxiety, while other patients were considered as not suffering from anxiety. Two thousand four hundred seventy-nine patients were included in this analysis.

### Metabolomics

Non-targeted global metabolomic profiling of serum derived from the BipOff cohort was performed by Metabolon (Durham, NC, USA) as previously described^112,113^. Briefly, samples were homogenized and subjected to methanol extraction then split into aliquots for analysis by ultrahigh performance liquid chromatography/mass spectrometry (UHPLC/MS) in the positive (two methods) and negative (two methods) mode. Metabolites were then identified by automated comparison of ion features to a reference library of chemical standards followed by visual inspection for quality control^114^. For statistical analyses and data display, any missing values are assumed to be below the limits of detection; these values were imputed with the compound minimum (minimum value imputation). Statistical tests were performed in ArrayStudio (Omicsoft) or “R” to compare data between experimental groups; p < 0.05 is considered significant and 0.05 < p 0.10 to be trending. An estimate of the false discovery rate (Q-value) was also calculated to take into account the multiple comparisons that normally occur in metabolomic-based studies, with q < 0.05 used as an indication of high confidence in a result.

*Targeted LPA quantification:* Mouse sera (15 µL) were extracted with 75 µL of ice-cold isopropanol (IPA) spiked with C17 cLPA as the internal standard (IS). To promote the lipid extraction and protein precipitation this solution was vortexed and centrifuged (at 4°C for 15 minutes at 14000rpm). The resulting supernatant was analyzed by reversed phase Liquid chromatography coupled to tandem mass spectrometry (RPLC-MS/MS) in negative ionization mode operating in selective reaction monitoring mode (SRM, TSQ Altis triple quadrupole system interfaced with a Vanquish UHPLC system (Thermo Fisher Scientific)). Chromatographic separation was carried out on a Zorbax Eclipse Plus C18 (1.8 μm, 100 mm × 2.1 mm I.D.) column (Agilent technologies, USA). Mobile phase was composed of A = 60:40 (v/v Acetonitrile: water solution) with 10 mM ammonium acetate and 0.1% acetic acid and B = 88:10:2 (Isopropanol: acetonitrile: water solution) with 10 mM ammonium acetate and 0.1% acetic acid. The linear gradient elution from 15% to 30% B was applied for 2 minutes, then from 30% to 48% B for 0.5 minutes, from 48% to 72% B and last gradient step from 72% to 99% B followed by 0.5 minutes isocratic conditions and a 3 min re-equilibration to the initial chromatographic conditions. The flow rate was 600 μL/min, column temperature 60 °C and sample injection volume 2µl. Data were processed using XCalibur 4.3 (Thermo Fisher Scientific) and concentrations were calculated to IS using a calibration curve.

### Genetic analyses of the PsyMetab cohort

All participants were genotyped on the Global Screening Array (GSA) v2 with multiple disease option (MD) chip at iGE3 institute in Geneva, Switzerland (http://www.ige3.unige.ch/genomics-platform.php). All quality control (QC) and filtering steps were performed in PLINK^115^. Ancestry was determined using Snpweights, a software for inferring genome-wide (GW) ancestry using SNP weights precomputed from large external reference panels^116^. Only individuals from European ancestry were considered in the present study. Multivariable logistic mixed models adjusting for age, sex, smoking and body mass index were conducted to test associations between SNPs and the anxiety phenotype. A dominant transmission was considered for rs10817103C>T (i.e., patients carrying the T allele were compared to patients carrying the CC genotype). Statistical significance was defined as a p value ≤ 0.05. Statistical analyses were performed using Stata 14 (StataCorp, College Station TX, USA) and R environment for statistical computing version 4.1.1.

### Single-cell RNA sequencing of human hippocampus

For the dataset of Ayhan et al., 2021^58^, count matrix was downloaded from GEO under accession number GSE160189. For the dataset of Tran et al. 2021^59^, count matrix and annotated metadata was downloaded from Github (https://github.com/LieberInstitute/10xPilot_snRNAseq-human).

*Cell filtering and quality controls:* To filter only high-quality cells using similar quality control criteria among the two databases, we applied filters on unique molecular identifier (UMI), mitochondrial and genes expressed counts per cell. We first filtered cells based on the percentage of UMI associated with a maximum of 12% mitochondrial transcripts expression. We further excluded cells with UMI and genes numbers above 3 MADs of the population median, with a minimum threshold defined at 100 detected genes. Potential doublets were removed using Scrublet^117^.

*Data integration and visualization:* To generate a reference human hippocampus database, we applied data integration procedure from Seurat and identified shared sources of variation between Ayhan et al. and Tran et al. datasets. Briefly, each dataset was normalized and scaled using SCTransform procedure from Seurat. We then identified common features and used the “FindIntegrationAnchors” function using for normalization method “SCT” and default parameters. We then performed the integration using the “IntegrateData” function and default parameters. For UMAP visualization, dimensionality reduction was performed using standard function in Seurat. We then use the Transfer Data function from Seurat to automatically annotate all the cells based on the reference annotations of Tran et al. dataset.

### Statistical analysis

All values are given as mean ± S.E.M. For the Fig. 1, results obtained in the validation of the BBA assay experiments were all analyzed by a one-way analysis of variance (ANOVA), with serum concentration as fixed factor. Analyses were followed by the Bonferroni post hoc test when appropriate. Results obtained in the BBA assay (*in vitro* proliferation) and for adult neurogenesis *in vivo* (quantification for Ki-67/BrdU and DCX) were all analyzed by an unpaired Student t-test. Results obtained in basal anxiety behavior experiments (EPM, LDT and anxiety scores) were all analyzed by an unpaired Student t-test. In Supplementary Fig. 1, results obtained from cell viability experiments (nuclear condensation and C-CASP3 activation) were all analyzed by the non-parametric Kruskal-Wallis test followed by a multiple comparison, with serum as fixed factor. Results obtained in chronic restraint stress experiment (LDT, EPM, anxiety scores and Free-corticosterone analysis) were all analyzed by an unpaired Student t-test. Body weight evolution was analyzed by a two-way analysis of variance (ANOVA) with repeated measures, with stress and days as fixed factors. Results obtained after group segregation (HP vs. LP and resilient vs. susceptible) were analyzed by the non-parametric Kruskal-Wallis test followed by a multiple comparison. In Fig. 2, results obtained in the human study (in vitro proliferation in the BBA assay, trait- and state-anxiety, and metabolites) were all analyzed by a one-way analysis of variance (ANOVA), with psychiatric disorders as fixed factors. Analyses were followed by Tukey’s multiple comparisons post hoc test when appropriate. Results obtained from in vitro cLPA16:0 treatment was analyzed by a one-way analysis of variance (ANOVA), with cLPA16:0 as fixed factor. Analysis was followed by the Bonferroni post hoc test. The BBA assay data with LPA_1_ antagonist treatment was analyzed by an unpaired Student t-test. Results obtained in cell viability in human study were analyzed by the non-parametric Wilcoxon-Mann-Whitney test. In Fig. 2A-T and Fig, 3A-W, results obtained were all analyzed by an unpaired Student t-test and one sample t-test with theoretical mean for anxiety score set at 0.5. Correlations were all analyzed by a Pearson correlation coefficient (r) calculation to establish relationships between dependent variables. In Fig. 3X-β, results obtained were all analyzed by a one-way analysis of variance (ANOVA), with drug treatment as fixed factors. Analyses were followed by Tukey’s multiple comparisons post hoc test when appropriate. All statistical tests were performed with GraphPad Prism (GraphPad 9 software, San Diego, CA, USA) using a critical probability of p < 0.05. Statistical analyses performed for each experiment are summarized in each figure legend with the chosen statistical test, sample size ‘n’ and p values, as well as degree of freedom and F/t values.

## Acknowledgments

The authors wish to thank Rebecca Borreggine and Tony Teav for technical assistance with mass spectrometry analyses, Fulvio Magara for help with animal behavioral experiments and the Cellular Imaging Facility of the University of Lausanne, for their assistance with microscopy.

## Funding

Swiss National Science Foundation 32003B_156914 (CBE)

Swiss National Science Foundation 31003A_173128 (CP)

Swiss National Science Foundation 310030_201015 (NT)

Swiss National Center of Competence in Research; “Synapsy: the Synaptic Basis of Mental Diseases” 51NF40-185897 (PM).

ERC starting grant CERDEV_759112 (LT)

## Author contributions

Conceptualization: NT, CP, AD, TL

Methodology: TL, CC, FG, CW, KG, AD, HGA, MT, HAC, JI, LT

Investigation: TL, CC, FG, AD, HAC, MT

Funding acquisition: NT, CP, AD, PM, CBE

Supervision: NT, CBE, HAC, JL, CP

Writing: TL, NT, AD, CBE, JI, LT, CP, HAC, CBE, PM, CP

