## Supplementary Figures for "Platelet-derived LPA16:0 inhibits adult neurogenesis and stress resilience in anxiety disorder"

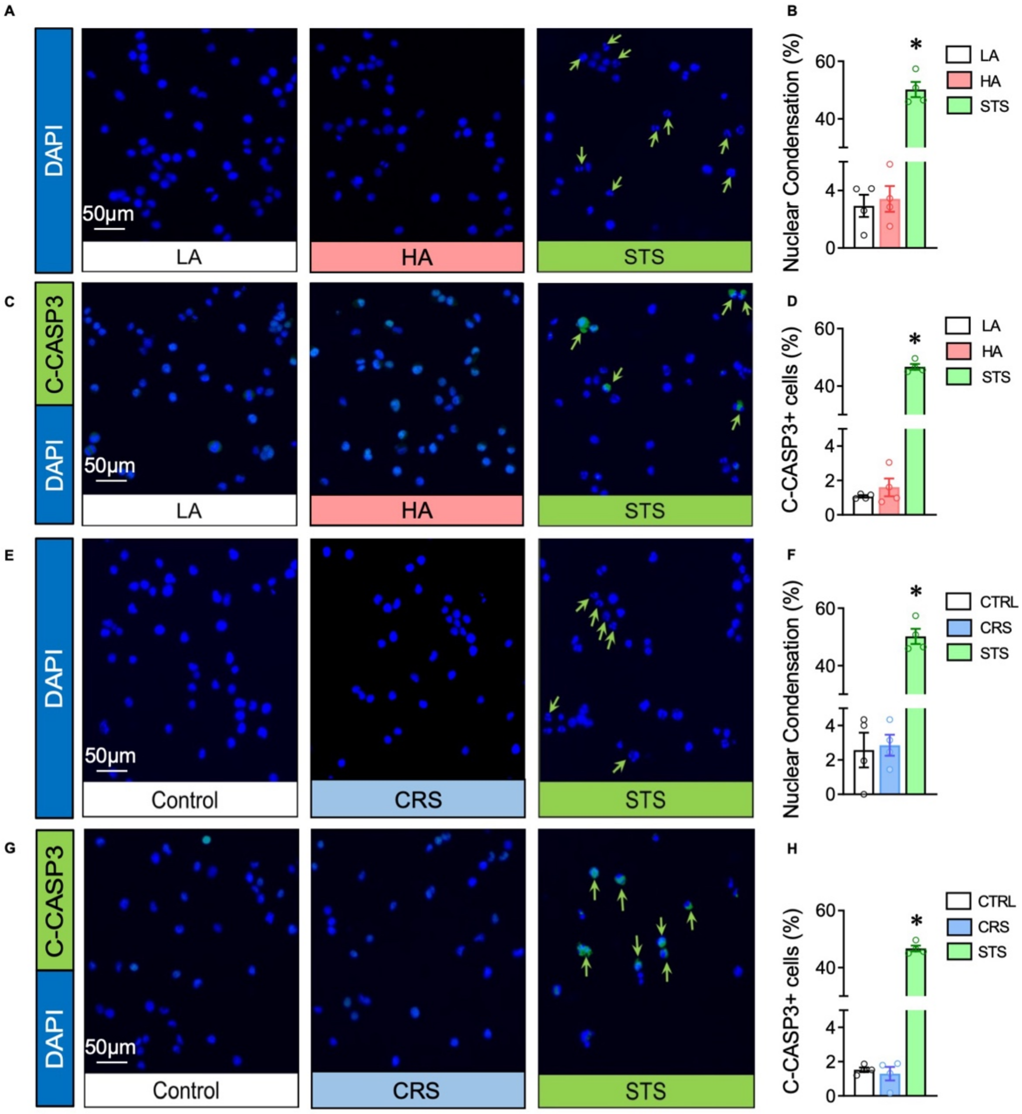


### Supp Fig. 1

**Supplementary Fig. 1: Serum from anxious mice does not affect cell viability. A**. Micrographs of aNPC exposed to serum from LA or HA mice (0.2% for 24h) or staurosporine (STS, 0.5 µM for 6h.) as positive control. Arrows point to condensed nuclei. **B**. Proportion of condensed nuclei in aNPC (Kruskal-Wallis test, $p = 0.0132$; multiple comparisons LA vs. HA: $p > 0.05$, LA vs. STS: $p = 0.028$; n = 3-4 per group). **C**. Micrographs of aNPC exposed to serum from LA or HA mice, or STS and immunostained for cleaved CASP3 (c-CASP3, green). Arrows point to c-CASP3$^+$ cells. **D**. Proportion of cells expressing c-CASP3 (Kruskal-Wallis test, $p = 0.0107$; multiple comparisons with the mean of LA group, LA vs. HA: $p > 0.05$, LA vs. STS: $p = 0.022$; n = 3-4 per group). **E**. Micrographs of aNPC exposed to serum from control mice, from CRS mice or to STA. Arrows point to condensed nuclei. **F**. Proportion of condensed nuclei in aNPC (Kruskal-Wallis test, $p = 0.0135$; multiple comparisons with the mean of CTRL group, CTRL vs. CRS: $p > 0.05$, CTRL vs. STS: $p = 0.032$; n = 3-4 per group). **G**. Micrographs of aNPC exposed to serum from control mice, CRS mice or to STS and immunostained for c-CASP3 (green). Arrows point to c-CASP3$^+$ cells. **H**. Proportion of c-CASP3$^+$ cells (Kruskal-Wallis test, $p = 0.0145$; multiple comparisons with the mean of CTRL group, CTRL vs. CRS: $p > 0.05$, CTRL vs. STS: $p = 0.05$; n = 3-4 per group). Histograms show average ± SEM. * $p<0.05$; ** $p<0.01$; *** $p<0.001$; ns: not significant.

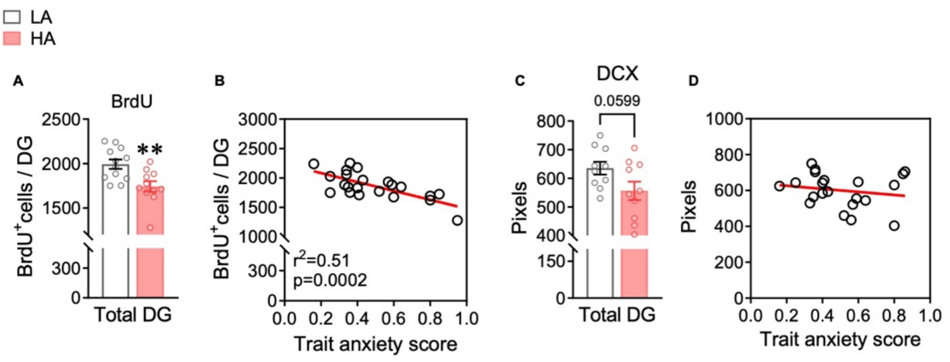


### Supp Fig. 2

**Supplementary Fig. 2: Trait anxiety is associated with decreased cell proliferation in the DG. A**. Histogram showing BrdU$^+$ cells in the DG expressed as the percentage of LA animals (DG, total: t20 = 2,470, p < 0.01, unpaired t-test, two-tailed, n = 11 per group). **B**. Correlation plot between trait anxiety scores and cell proliferation in the DG. **C**. Histograms of the proportion of DCX immunostained pixels in the DG (t19 = 2.01, p = 0.0599, unpaired t-test, two-tailed, n = 10-11 animals per group) of LA (white) and HA (red) mice. **D**. Correlation plot between trait anxiety score and DCX immunostaining in the DG. Linear regression, $r^2$ and p values are indicated in the graphs when significant correlations were found. * p<0.05; ** p<0.01; ns: not significant.

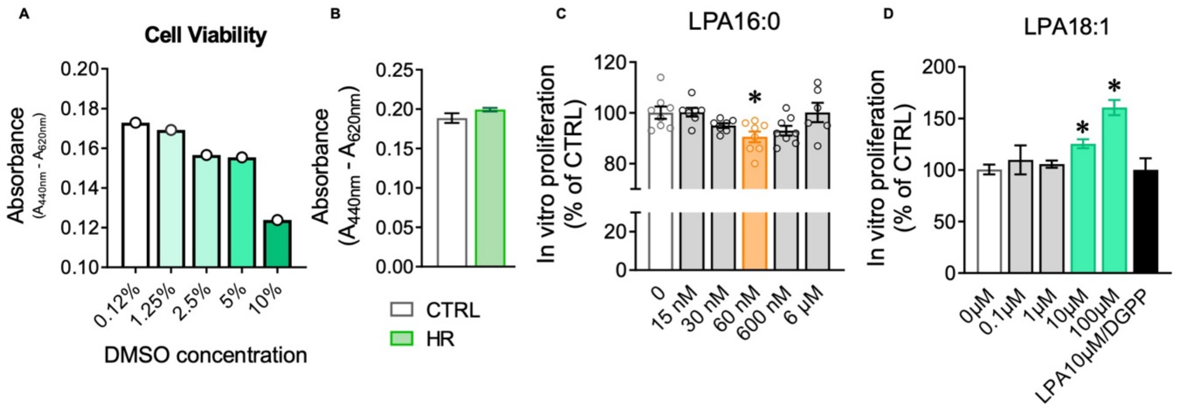


### Supp Fig. 3

**Supplementary Fig. 3: Serum from HR patients does not affect cell viability. Opposite effects of LPA16:0 and LPA18:1 on aNPC proliferation. A.** Dose-response effect of DMSO (as positive control) on cell viability, as measured by absorbance. **B.** Effect of serum from CTRL and HR subjects on cell viability. **C-D**. Dose-response effect on aNPC proliferation of **(C)** LPA16:0 treatment ($F_{4,38}$ = 3.744, p = 0.0075, one-way ANOVA; n = 4 wells per condition for each replicate, 2 independent replicates) and **(D)** LPA18:1 treatment (p = 0.0113, Kruskal-Wallis test, post-hoc analysis for multiple comparisons: 0µM vs. 10µM, p = 0.05; 0µM vs. 100µM, p = 0.029). * p<0.05.

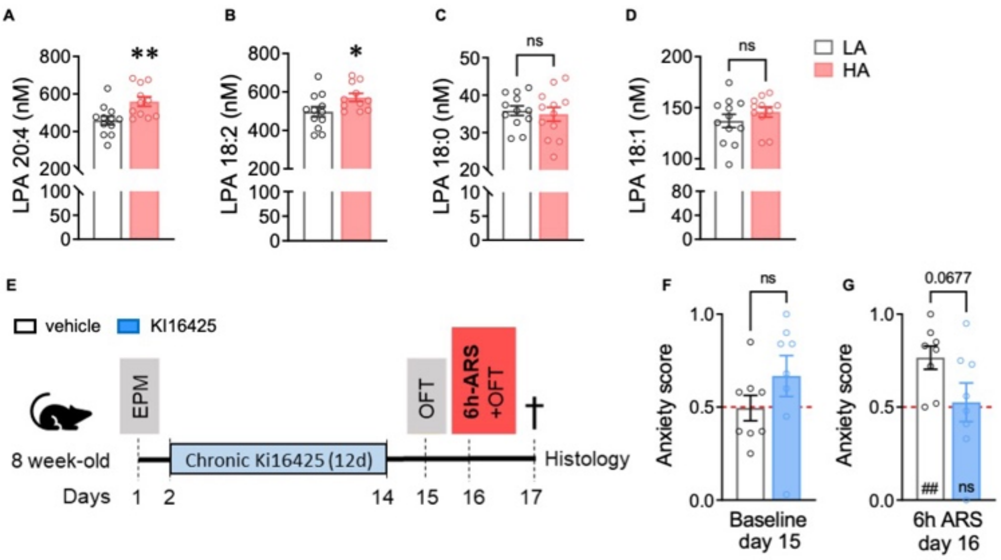


### Supp Fig. 4

**Supplementary Fig. 4: Serum concentrations of different forms of LPA in anxious mice and replication study for the effect of Ki16425 on resilience to a 6h ARS. A-D** Histograms of the concentration of LPA20:4 (**A**), LPA18:2 (**B**), LPA18:0 (**C**) and LPA18:1 (**D**) in the serum of LA vs HA mice (LPA20:4; $t_{21} = 2.968$, $p = 0.0073$, LPA18:2; $t_{21} = 2.206$, $p = 0.0387$; LPA 18:0; $t_{21} = 0.4188$, $p > 0.05$, LPA 18:1; $t_{21} = 1.060$, $p > 0.05$, unpaired t-test, two-tailed, n = 11 per group). **E.** Experimental design for *in vivo* Ki16425 administration. **F, G.** Histograms of the anxiety score of mice injected with vehicle (white) or Ki16425 (blue) after 12 days of treatment under **(F)** baseline conditions ($t_{14} = 1.329$, $p > 0.05$, unpaired t-test, two-tailed; vehicle: $t = 0.07324$, $p > 0.05$, One sample t test and Ki16425: $t = 1.517$, $p > 0.05$, One sample t test, n = 8 mice per group) and **(G)** after 6 hours of ARS ($t_{14} = 1.980$, $p = 0.0677$, unpaired t-test, two-tailed; vehicle: $t = 4.256$, $p = 0.0038$, One sample t test and Ki16425: $t = 0.2529$, $p > 0.05$, One sample t test, n = 8 mice per group). Histograms show average ± SEM. * $p<0.05$; ** $p<0.01$; ns: not significant. Comparison between the group mean and the hypothetical value of 0.5, to assess anxiety withing each group are shown within each histogram bar. One-sample t-test. #: $p<0.05$; ##: $p<0.01$.
